# scPRIME-seq uncovers synchronized multimodal molecular collapse facilitated by pre-neoplastic epigenetic burden in prostate cancer

**DOI:** 10.64898/2026.10.07.757464

**Authors:** Shengdi Li, Ziqun Liu, Yingying Xie, Xinan Wang, Hanchu Ye, Xingyong Ma, Ming Tan, Denglong Wu, Shengsong Huang, Zhenfei Li

## Abstract

Tumorigenesis involves alterations across the genome, epigenome and transcriptome, yet how these layers interact to drive malignant transformation remains unclear, largely due to the lack of scalable multimodal single-cell approaches. Here, we developed scPRIME-seq, a high-throughput platform for simultaneous single-cell profiling of transcriptome, genome and DNA methylome, and applied it to 436,908 cells from 29 prostate cancers and 15 benign tissues. This multimodal atlas resolved dynamic changes across 30 cell types and a unified tumorigenic trajectory. Integrated analysis revealed a cell-state transition “tipping point”, where the onset of aneuploidy is temporally coupled with global methylomic alterations and transcriptomic rewiring. Early focal epigenetic alterations, including *TMEM106A* promoter hypermethylation and the emergence of hypomethylated domains, were present in founder aneuploid clones and benign tissues, indicating a pre-neoplastic origin of epigenetic burden. These pre-neoplastic alterations coincide with transcriptomic changes that affect epithelial interactions with stromal and immune compartments, promoting a putative tumor-permissive microenvironment. scPRIME-seq provides a high-throughput multimodal framework for identifying conserved, cell-intrinsic molecular programs underlying tumorigenesis.

## Introduction

Elucidating the precise regulatory mechanisms that govern cellular identity and state transitions demands integrated interrogation of the genome, epigenome, and transcriptome. The necessity for such multimodal profiling is particularly compelling in cancer research, where diverse DNA methylation alterations occur across tumor types and disease stages^1^, including widespread hypomethylation in gene-poor, heterochromatic regions and focal hypermethylation at CpG island-associated loci^2^. In prostate cancer (PCa), extensive DNA mutations accumulate predominantly in metastatic disease, while remaining rare in early-stage primary tumors, suggesting that epigenetic plasticity may be a major driver of PCa initiation and malignant transition^3,4^. Dysregulation of DNA methylation has been documented across different stages of PCa^5,6^, with aberrantly methylated genomic domains or genes serving as candidate biomarkers for disease onset and progression^7–9^.

Distinguishing molecular alterations that drive stable cell-state transitions from transient cellular responses or stochastic variation is essential for uncovering disease-driving mechanisms. While transcriptome-based studies have revealed extensive inter- and intratumoral heterogeneity^10^, gene expression states are shaped by diverse influences, including genetic lesions, epigenetic remodeling, lineage history and microenvironmental stresses^11–13^, as well as stochastic variability and sample-preparation effects that can artificially amplify transient stress responses^14,15^. Transcriptomic variation alone therefore cannot readily determine whether a given cell state is anchored by stable genomic or epigenomic alterations^16,17^. Conversely, cancer genomes and epigenomes accumulate widespread alterations, many of which may not lead to recurrent phenotypic consequences. Thus, elucidating the molecular basis of tumorigenesis requires single-cell technologies that can link upstream genomic and epigenomic alterations to downstream transcriptomic changes. This capability is particularly important for understanding cell state reprogramming during tumorigenesis, which is believed to arise through a combination of intrinsic genetic and epigenetic changes, and microenvironmental influences^18^. A recent colorectal cancer study identified the emergence of a metastatic cell state in the earliest stage of tumorigenesis, suggesting that such state transitions may occur as early events rather than late consequences of metastatic progression^19^. Consequently, robust high-throughput multimodal single-cell platforms are essential for resolving rare cell populations within highly heterogeneous tumors.

Despite recent advances enabling simultaneous profiling of the single-cell DNA methylome and transcriptome, existing approaches largely rely on plate-based isolation of individual cells or nuclei (e.g., scTrio-seq2, scNMT-seq)^17,20^, due to the requirement for cell lysis and physical partitioning of DNA and RNA prior to cell barcoding. This conventional workflow typically limits throughput to hundreds of cells per sample, restricting its application in highly heterogeneous systems such as tumors, where large cell numbers are required to resolve rare populations and transitional states. Alternative strategies combine single-nucleus methylation sequencing with scRNA-seq on separately prepared samples to approximate joint profiling, but lack direct molecular linkage between the two modalities^21^. A scalable method capable of profiling tens to hundreds of thousands of cells would enable the reconstruction of dynamic epigenetic and transcriptomic trajectories during disease progression, yet such an approach remains lacking.

Here, we introduce <u>P</u>ermeabilization-based <u>R</u>NA Integrated DNA <u>ME</u>thylation sequencing (scPRIME-seq), an ultra-high-throughput single-cell technology for parallel and scalable profiling of the DNA methylome, genomic copy number variations (CNVs), and 5’ transcriptome. Applied to 44 patient-derived samples, comprising a multi-omic dataset of 436,908 cells, scPRIME-seq enables construction of a comprehensive single-cell atlas of primary prostate cancer. Our study reveals a cell-state transition “tipping point” during disease onset with waves of multimodal molecular alterations and delineates their temporal relationships, thereby highlighting the biological and clinical potential of multimodal single-cell technologies for cancer diagnosis and research.

## Results

### Development of scPRIME-seq workflow

scPRIME-seq is designed to enable scalable joint profiling of transcriptomes and DNA methylomes in single cells by overcoming the limited compatibility between droplet-based workflows and the concurrent capture of RNA and DNA. A central challenge was to achieve efficient nuclear access by Tn5 transposase through permeabilized cell membrane and disrupted nucleosome structure, while maintaining cytoplasmic and nuclear RNA integrity.

To address this, we developed an optimized methanol-based cell permeabilization and *in situ* tagmentation protocol that simultaneously satisfies these requirements. This approach supports uniform Tn5 transposition across genomic DNA within intact cells, as indicated by substantially reduced positional bias towards open chromatin regions (**Supplementary Fig. 1a**). Unlike formaldehyde crosslinking employed in sciMET and comparable platforms^22,23^, methanol fixation preserves RNA integrity in both the cytoplasm and nucleus^24,25^, thereby enabling efficient 5’ labeling of mRNA via reverse transcription (RT) and template switching (**Fig. 1a**). Consistently, no detectable loss of RNA was observed following treatment (**Supplementary Fig. 1b-c**). By obviating the need for pre-barcoding physical partitioning of RNA and DNA fractions, scPRIME-seq removes a key throughput bottleneck in conventional dual-omic single-cell methods (e.g., scTrio-seq2, scM&T-seq)^17,26^, thereby enabling fully droplet-compatible multimodal profiling and substantially improving scalability (**Fig. 1a**).

**Figure 1.**
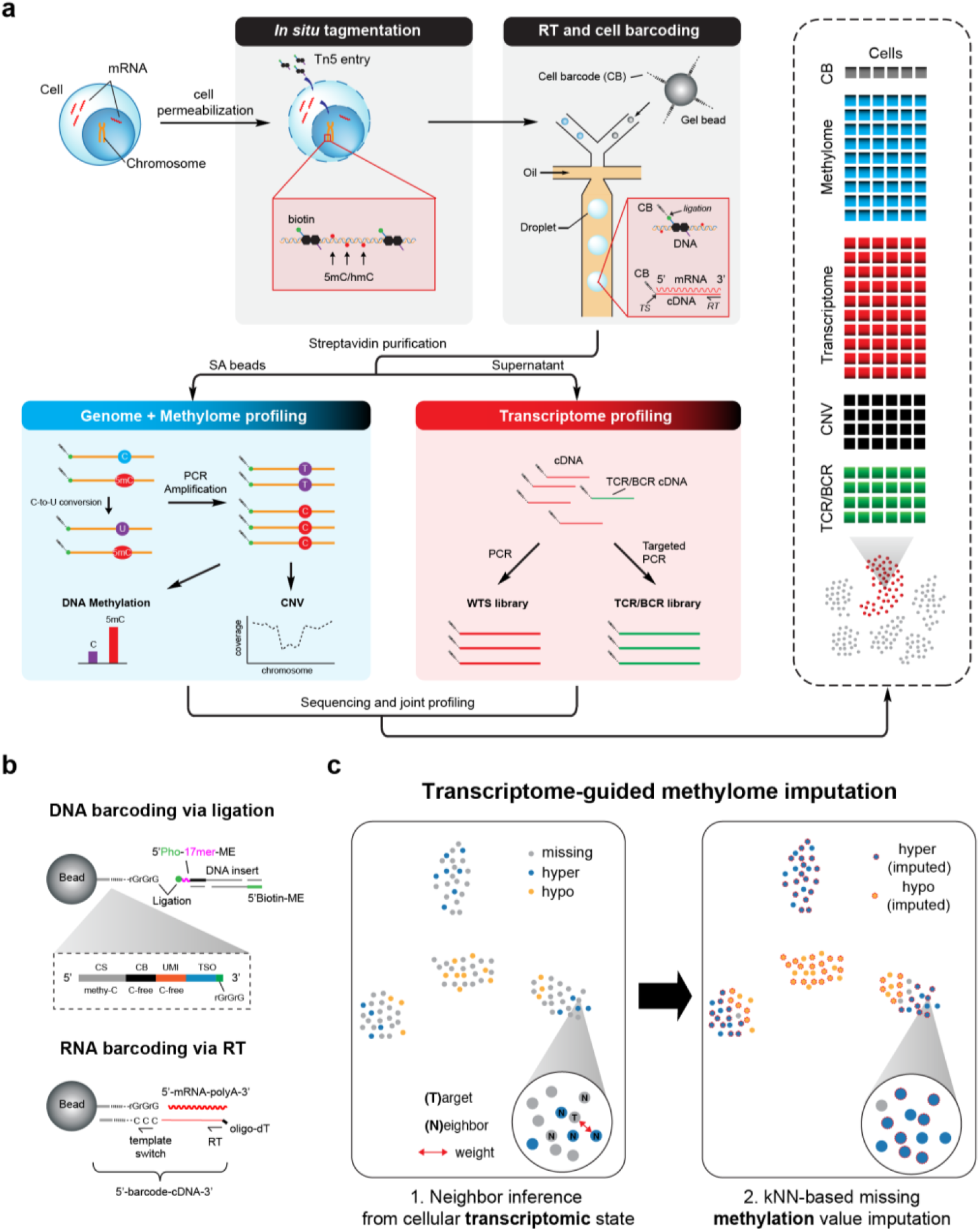
A scalable workflow for joint profiling of single-cell transcriptomes and DNA methylomes. **a**, Schematic illustration of scPRIME-seq workflow. **b**, Design of a dual-purpose bead-conjugated cell barcode oligonucleotide. The 3′ end of the single-stranded DNA barcode oligo contains three riboguanosines (rGrGrG), which function as a template-switching oligo (TSO) by annealing to the non-templated 5′-CCC overhang at the 5′ end of cDNA to enable incorporation of the 5′-labeled cell barcode. This ribonucleotide-containing terminus also facilitates ligation to 5′-phosphorylated transposed DNA fragments via T4 RNA ligase 2. **c**, Pipeline for imputing incomplete single-cell methylomes using k-nearest neighbors (kNN) of transcriptomically similar cells. CB: cell barcode; CNV: copy number variation; ME: mosaic end; RT: reverse transcription; TS: template switch; TSO: template switch oligo; CS: constant sequence; WTS: whole-transcriptome sequencing; TCR/BCR: T-/B-cell receptors.

Another key feature of scPRIME-seq is a custom-designed bead-conjugated cell barcode oligonucleotide that enables simultaneous labeling of genomic DNA and 5’ RNA from the same cell. The oligonucleotide terminates with three riboguanosines (rGrGrG) at its 3’ end, enabling template switching to append cell barcodes to the 5’ end of full-length cDNA (**Fig. 1b**). This design enables 5’ transcriptome sequencing, including the variable region of immune receptor V(D)J sequences for profiling immune cell populations. Moreover, the 3’ rGrGrG end also supports ligation to transposed genomic DNA fragments bearing a 5’ phosphate using T4 RNA ligase 2 (**Supplementary Fig. 1d**). Cytosine-free cell barcodes and unique molecular identifiers (UMIs) were used to ensure compatibility with enzymatic C-to-U conversion during methylome processing.

We also redesigned the transposase system to ensure compatibility with the workflow. Specifically, the leading sequence upstream of the mosaic end (ME) was modified to a 17-mer bearing a 5’ phosphate to facilitate barcode ligation (**Fig. 1b**). In parallel, a gap-filling step incorporating methylated dCTP was introduced, generating a chimeric ME region flanking each captured DNA fragment with strand-specific cytosine methylation (**Supplementary Fig. 1e**; **Materials and Methods**). This region serves as an internal readout for read-level conversion efficiency based on strand-specific C-to-T conversion rates. Following droplet-based barcoding and RT, biotin-labeled genomic DNA (introduced via biotin-Tn5) was separated from cDNA, enabling parallel profiling of DNA methylome and transcriptome from the same cell. The RNA fraction was subsequently processed using a 5’-based sequencing strategy, enabling recovery of T cell receptor (TCR) and B cell receptor (BCR) sequences alongside whole-transcriptome profiles (**Supplementary Fig. 1e**). In benchmarking experiments using five peripheral blood mononuclear cell (PBMC) samples (3,000 cells each), scPRIME-seq achieved an average dual-omic capture rate of 68.7% (1,781-2,561 cells per sample) and exhibited consistent transcriptome data quality (median 1,307 genes per cell) and stable cell-type compositions, demonstrating reproducibility of the method (**Supplementary Table 1**).

To overcome the intrinsic sparsity of single-cell DNA methylation data, we developed a computational framework, named sc<u>PRIME</u>-seq compatible Inference of <u>M</u>ethylation <u>P</u>rofiles <u>U</u>sing <u>T</u>ranscriptomic <u>E</u>vidence (PRIME-IMPUTE), that leverages transcriptomic similarity to reconstruct methylomes (**Fig. 1c**). PRIME-IMPUTE assumes that transcriptomically similar cells share broadly similar DNA methylation patterns. It increased effective methylome coverage on a test dataset (RS67) from 16.6% to 92.7%, with an imputation accuracy of 70.2%, while preserving cell-type-specific methylation patterns (**Supplementary Fig. 2**).

Together, scPRIME-seq provides a scalable and efficient solution for high-throughput joint profiling of gene expression and DNA methylation at the single-cell level, with improved chromatin accessibility, reduced experimental complexity, and enhanced data completeness compared to existing technologies.

### A single-cell transcriptome and DNA methylome atlas in prostate cancer

To delineate the dynamic molecular alterations during cancer initiation and malignant progression at the single-cell level, we applied scPRIME-seq to samples derived from 44 patients, stratified into three groups (G1, G2 and G3) according to disease stage (**Fig. 2a**). PCa groups (G2 and G3) were defined by Gleason score (GS), with GS < 8 as low- to intermediate-risk cancer (n = 11) and GS ≥ 8 as high-risk prostate cancer (n = 18). G1 specimens were histologically confirmed as benign tissues (n = 15), thereby serving as non-tumor controls in this study (**Supplementary Table 2**). Following quality control, we retained a total of 496,543 cell barcodes, with an average of 11,285 cells per sample. Among these, 436,908 cells were captured in both transcriptomic and methylomic profiling, representing 95.5% (436,908/458,484) and 91.8% (436,908/476,132) of barcodes recovered by each single modality, respectively.

**Figure 2.**
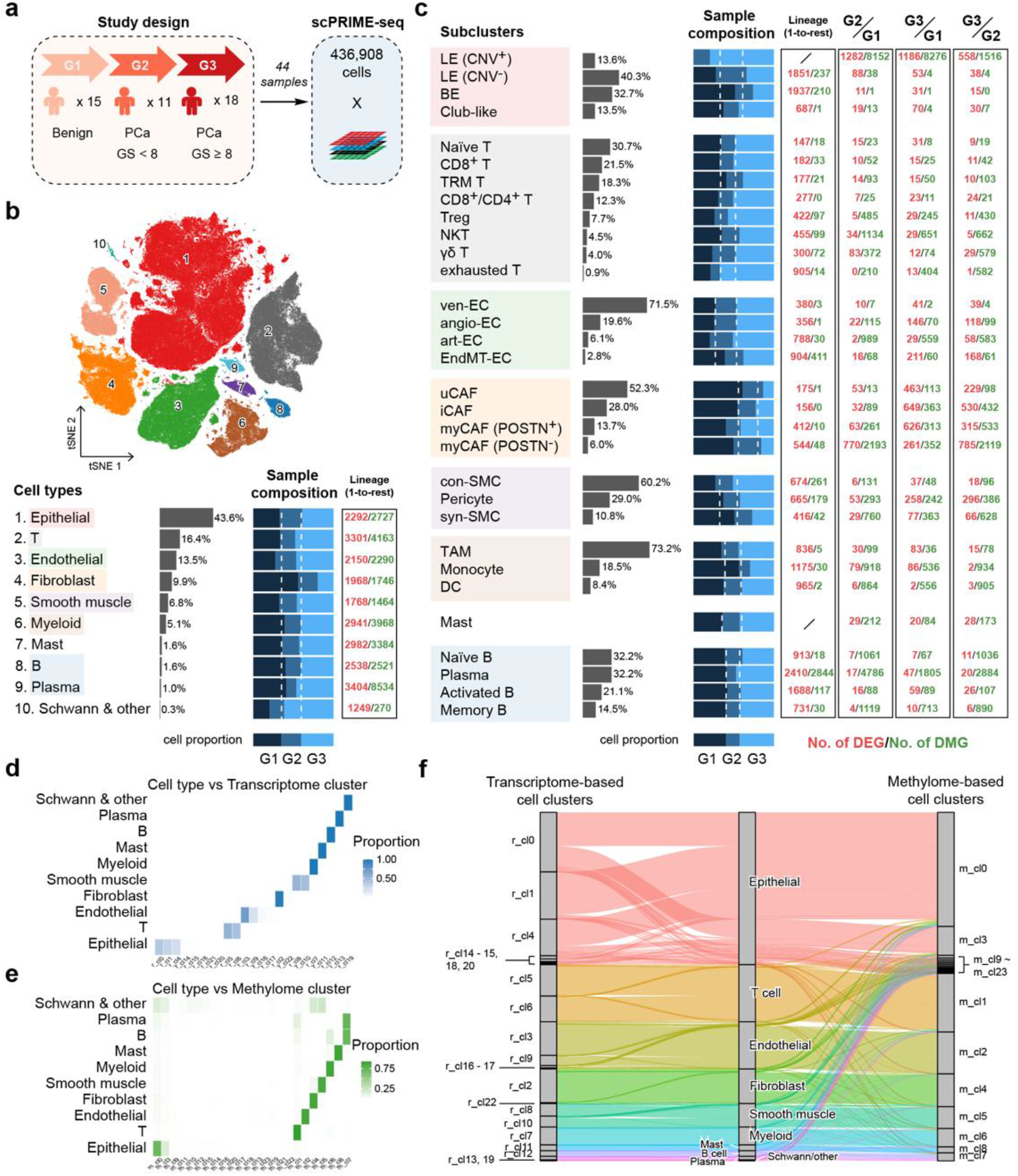
A single-cell transcriptome and DNA methylome atlas of prostate cancer. **a**, Grouping of patient-derived samples subjected to scPRIME-seq, including benign prostatic tissues and prostate cancer (PCa) stratified by Gleason score (GS). **b-c**, t-SNE visualization of 458,484 cells captured by transcriptomic sequencing, subdivided into major cell lineages (lower part of panel **b**), and further into subclusters by secondary clustering (panel **c**). White dashed lines represent the expected distribution of cell numbers based on the overall cell composition across the three sample groups G1-G3. The number of differentially expressed genes (DEGs) and promoter-associated differentially methylated genes (DMGs) are shown for one-vs-rest comparisons between cell types (“lineage”) and for between-group comparisons within each cell type. **d-e**, Cell type composition of Leiden clusters derived from transcriptomic (**d**) or methylomic (**e**) data. Color intensity represents the proportion of each cell type (rows) within each cell cluster (columns). **f**, Alluvial plot showing the correspondence between transcriptome-based clusters (left) and methylation-based clusters (right), linked to 10 annotated cell types (center).

Clustering analysis identified 10 major cell types and 30 subclusters across benign and malignant prostate samples and quantified their distribution across G1-G3 (**Fig. 2b-c**; **Supplementary Fig. 3-4**). Methylome-based clusters (without imputation) closely matched transcriptome-defined clusters, indicating that DNA methylation serves as a key determinant of cell lineage identity and is predictive of cell state (**Fig. 2d-f**; **Supplementary Fig. 5**). Across genomic regions, DNA methylation was generally higher within gene bodies, whereas promoter regions exhibited more variable methylation levels (**Supplementary Fig. 6a**). Gene expression showed a negative correlation with promoter methylation level (r = −0.23; **Supplementary Fig. 6b**), while gene body methylation showed little to no correlation with transcriptional activity (r = 0.002; **Supplementary Fig. 6c**). Across all cell types, hematopoietic lineages (T, B and plasma cells) exhibited globally lower levels of DNA methylation, whereas epithelial and stromal populations (endothelial, smooth muscle cells and fibroblasts) showed higher methylation levels (>80% on average). Mast and myeloid cells exhibited the most pronounced hypermethylation (**Supplementary Fig. 6d**).

A key strength of single-cell DNA methylation sequencing is that DNA fragment distributions can be used to infer chromosomal copy number variations (CNVs). Accordingly, integrating CNV information in scPRIME-seq enables more accurate and refined cell identity classification. For instance, the presence of CNVs allows classification of 13.6% of luminal epithelial (LE) cells as CNV^+^ tumor cells, which emerged and expanded in G2 and G3 samples (**Fig. 2c**), in contrast to CNV^−^ LE cells representing benign or transitional states without detectable genomic alterations. Furthermore, joint profiling of gene expression and DNA methylation improves the resolution of cell subtype identification, enabling the detection of previously unrecognized populations. For example, a T-cell-like population was identified by the absence of *CD3* and *TRAC* expression, but with the presence of *TRDC*, suggesting a potential γδ T lineage commitment without functional CD3 complex assembly. Hypermethylation of the *CD3* and *TRAC* promoters further rules out the possibility of *CD3* false-negative expression, providing solid evidence for the existence of this rare subcluster (**Supplementary Fig. 7**).

To evaluate the role of DNA methylation in cell lineage establishment, we analyzed differentially expressed genes (DEGs) and differentially methylated genes (DMGs; defined as genes with an adjusted *p* value < 0.05 and >15% promoter methylation change) (**Fig. 2c**). About 6% of DMGs showed corresponding inverse changes in gene expression (i.e., hypomethylated DMGs associated with upregulated DEGs, and vice versa; **Supplementary Fig. 8a-b**). Notably, this overlap increased to >40% when restricting the analysis to the actively expressed transcriptome (genes expressed in ≥10% of cells in at least one cell type; **Supplementary Fig. 8c-d**). This striking contrast suggests that DNA methylation changes at regulatory regions occur more broadly than those directly coupled to transcriptional output, indicating that additional regulatory layers modulate whether methylation changes ultimately translate into transcriptional activation or suppression.

### Genomic copy number aberrations during prostate cancer progression

Given the unique advantages of scPRIME-seq, we analyzed somatic CNV patterns across all 44 specimens and identified a total of 162 subclones, including one diploid subclone without detectable alterations in each sample (**Fig. 3a**). 96.8% of aneuploid cells were assigned to epithelial cells, with 94.2% specifically assigned to LE cells (**Fig. 3b-c** and **Supplementary Fig. 9a**), consistent with previous reports that primary prostate cancer is predominantly of luminal origin^27^. The remaining 3.2% of aneuploid cells likely reflect misassigned cell types or false-positive calls inherent to single-cell CNV inference.

**Figure 3.**
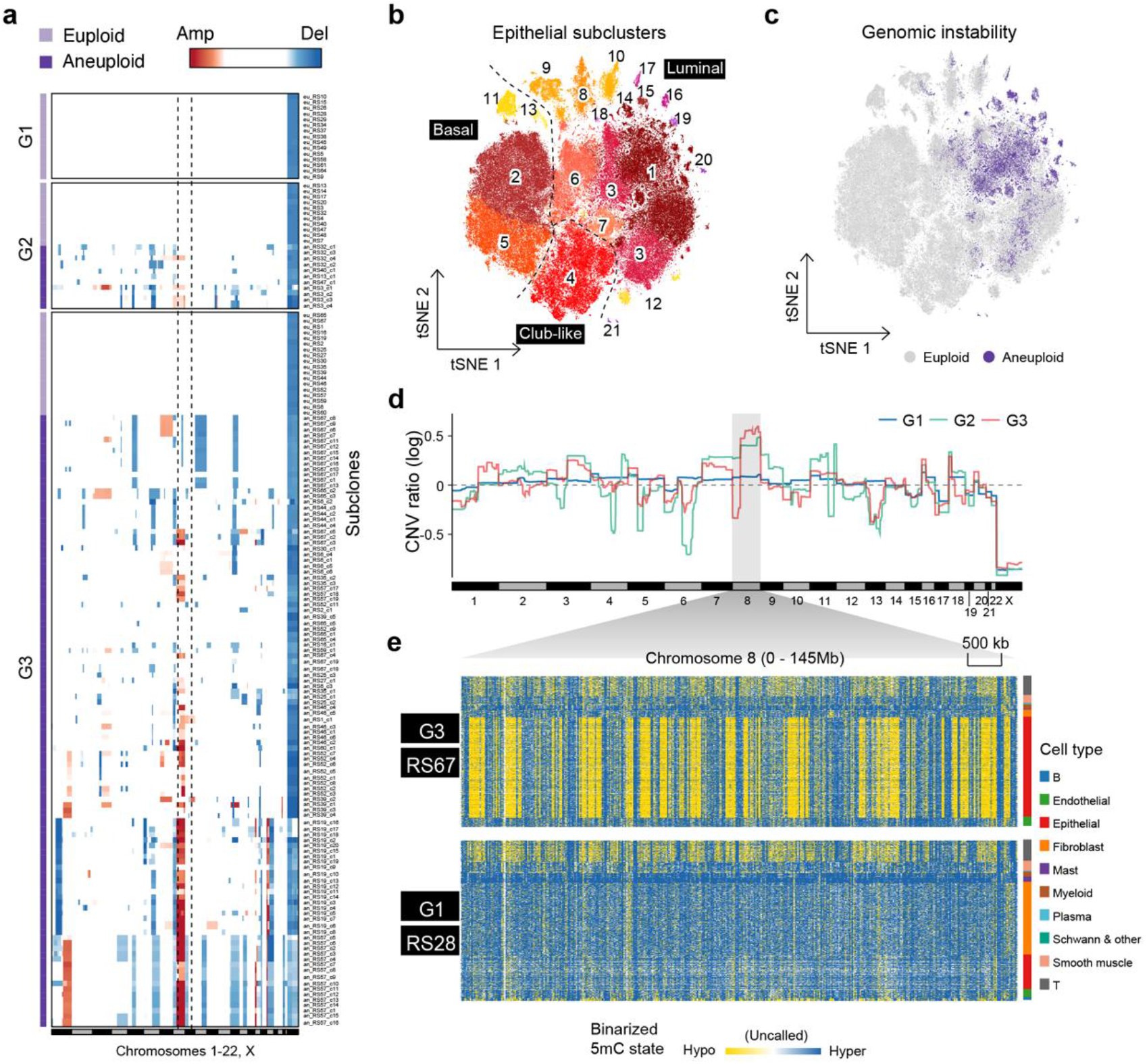
The genomic and epigenomic heterogeneity of prostate cancer cells. **a**, Distribution of CNVs across genomic regions and patient-derived subclones. Each row represents a subclonal karyotype shared by multiple cells within an individual patient. Only cells passing quality control for CNV inference were preserved in this analysis. **b**, t-SNE visualization of epithelial subclusters identified by secondary clustering of transcriptomic profiles within epithelial cells. **c**, CNV profiles overlaid onto t-SNE space in **b**. Cells showing gain or loss in any chromosomal arm are marked as aneuploid. **d**, Aggregated CNV ratio (log_2_-transformed) across all patient cells, stratified by G1-G3. A log_2_ ratio of 0 indicates the expected copy number of a diploid genome, with chromosome X representing a single-copy baseline in male samples. **e**, Heatmap visualization of DNA methylation status along chromosome 8 for a BPH sample (lower) and a PCa sample (upper). DNA methylation is shown as binarized values per 20-kb bin (yellow: hypomethylated, blue: hypermethylated, white: missing).

Among the three cohorts, no aneuploid LE cells were detected in G1, as expected, whereas G2 and G3 harbored 11 and 107 aneuploid subclones, respectively (**Fig. 3a**), suggesting a marked increase in intratumoral heterogeneity during disease progression. 5 out of 11 G2 patients showed no detectable CNVs, whereas all 18 G3 patients exhibited CNVs with variable aneuploid-to-euploid ratios (**Supplementary Table 2**). Subclonal size ranged from a few to >1000 cells, with no apparent difference between G2 and G3 (**Supplementary Fig. 9b**). However, given the much higher number of subclones (**Fig. 3a**; **Supplementary Fig. 9c**), the total aneuploid fraction and cell count were higher in G3 than in G2 (G2: 23.7%; G3: 59.5%), consistent with a more advanced disease stage.

The genomic distribution of amplification and deletion events revealed a high degree of concordance across subclones from different patients (**Fig. 3d**), consistent with recurrent driver CNVs previously reported in primary and metastatic prostate cancer^3,4^. For example, among 23 CNV-positive tumors, chromosome 8p loss was observed in 17 cases (73.9%), 8q gain in 14 (60.9%), chromosome 13 full or partial loss (encompassing *RB1* deletion) in 15 (65.2%), and 3q gain in 9 (39.1%) (Supplementary Table 3).

In aneuploid LE cells from PCa samples (G2 and G3), extensive hypomethylation within large, patterned genomic blocks termed partially methylated domains (PMDs)^7^ was evident, whereas this epigenetic signature was undetectable in non-tumor epithelium (G1) (**Fig. 3e**). However, the intrinsic and temporal relationships between PMD-associated hypomethylation and CNV alterations during cancer initiation and progression remain poorly defined. Notably, conventional bulk and unimodal single-cell sequencing approaches are unable to resolve the temporal relationships among these processes, which can now be interrogated using scPRIME-seq.

### A temporally coordinated multimodal molecular collapse defines a tipping point during prostate cancer initiation

To understand how benign LE cells progress towards genomic, epigenomic and transcriptomic collapse, we used the genome-wide mean hypomethylation fraction as a proxy for progression along global demethylation (**Materials and Methods**). This provided a continuous axis to map and interpret the trajectory of genomic, epigenomic and transcriptomic alterations during cancer progression.

Joint profiling of global DNA methylation and transcriptomic status on chromosome 8, a genomic region known to be among the earliest to acquire CNVs in PCa, revealed a highly synchronized pattern: decreased DNA methylation (**Fig. 4a**), increased aneuploid cell fraction (**Fig. 4b**), and drastic transcriptomic shifts (**Fig. 4c**) occurred concurrently. Within the same clinical specimen, the luminal compartment typically comprises a mixture of benign and tumor cells, with limited cells residing in an intermediate state (**Fig. 4d**). This intratumoral bimodal distribution of DNA methylation levels indicates the presence of a putative transition point in the benign-to-tumor transition, marked by rapid accumulation of molecular alterations and selective advantage, rendering intermediate states largely latent.

**Figure 4.**
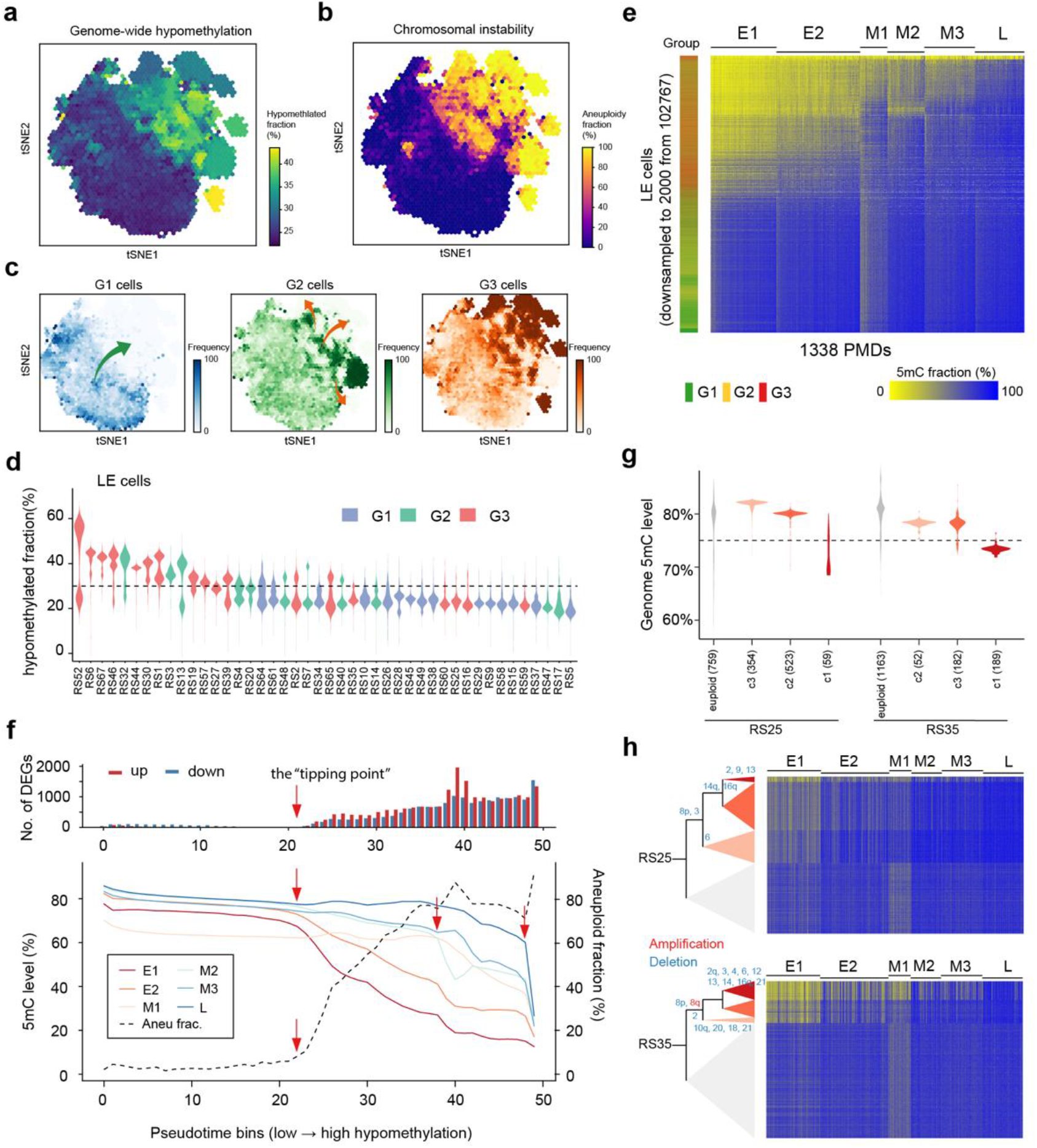
Concurrent onset of global hypomethylation, copy number aberration and transcriptomic rewiring. **a-c**, t-SNE embedding of LE cells based on transcriptomic profiles of genes located on chromosome 8, overlaid with genome-wide hypomethylation progression **(a**), increasing chromosomal instability (**b**) and the distribution of cells across G1-G3 groups (**c**). **d**, Distribution of genome-wide hypomethylation fractions per cell across LE cells for each patient. **e**, DNA methylation levels across 1,338 PMDs in LE cells from all patients. For visualization and clustering efficiency, LE cells were randomly downsampled to 2,000 cells. Cells were ordered by genome-wide hypomethylation fraction (top, high; bottom, low), and PMD clusters were defined by hierarchical clustering. **f**, A total of 102,767 LE cells were ordered by increasing hypomethylation fraction and partitioned into 50 bins of equal cell number (*x*-axis), representing a pseudotemporal trajectory of progressive genome demethylation. For each bin, PMD methylation levels and aneuploid fractions are shown. Differential gene expression per bin was assessed relative to all euploid LE cells. **g**, Genome-wide methylation of subclones from two prostate cancer patients (RS25, RS35) calculated as the mean CpG methylation fraction per cell. The number of cells within each subclone is indicated in parentheses. **h**, Subclonal evolution inferred from CNV patterns. Individual CNV events and their temporal order during intra-tumor evolution were determined by comparing CNV distribution across subclones within each patient.

By analyzing the genomic distribution of hypomethylated regions in epithelial cells, we identified a total of 1,338 PMDs across all samples (**Supplementary Table 4**). PMDs were absent in benign luminal cells but frequently and concurrently distributed in samples showing global hypomethylation (**Supplementary Fig. 10**). By reordering all 102,767 LE cells according to their genome-wide hypomethylation fraction, we reconstructed a pseudotemporal trajectory of epigenomic evolution in prostate cancer. This trajectory revealed a clear progression from G1 to G2 and G3 cells (**Fig. 4e**). The 1,338 PMDs were classified into 6 clusters based on their timing during tumor progression (E1, E2: early-stage PMDs; M1, M2, M3: mid-stage PMDs; L: late-stage PMDs). Interestingly, the initial rise in genome instability (measured as the fraction of aneuploid cells) was synchronized with the onset of DNA methylation loss in early PMDs at pseudotime bin 22 (**Fig. 4f**). Concurrent with this same point, extensive transcriptome rewiring also emerged, as indicated by a sharp increase in the number of differentially expressed genes relative to the euploid reference. We hereafter term this state transition point at bin 22 as the “tipping point”, which marks the onset of coordinated, extensive epigenetic, genomic and transcriptomic alterations during tumorigenesis (**Fig. 4f**).

Furthermore, a continuous progression comprising multiple phases of epigenomic degeneration was evident from the trajectory after the tipping point. In phase 1 (bin 22 - 37), gradual methylation loss occurs at E1 and E2 PMDs, accompanied by accumulation of CNVs. Phase 2 (bin 38 - 48) is characterized by relatively steady aneuploidy fraction at ∼75%, during which M1, M2 and M3 PMDs begin to demethylate. In the final phase 3, a sharp decrease of methylation is observed in L PMDs, together with a surge of aneuploidy fraction to ∼90% (**Fig. 4f**). These results demonstrate coordinated alterations across the epigenome, genome and transcriptome during tumor initiation and malignant progression, with such synergistic molecular perturbations originating from specific genomic regions.

To assess how multiomic evolution aligns with a transcriptome-centric trajectory, we reconstructed pseudotime using a standard transcriptome-based approach (**Materials and Methods**). While most normal and tumor cells form a continuum in UMAP space, transcriptome-based pseudotime does not define a clear normal-to-malignant transition point, underscoring a key limitation of unimodal analyses in tumor contexts (**Supplementary Fig. 11**). Moreover, tumor transcriptomes exhibit substantial inter-patient heterogeneity, leading to divergent evolutionary paths (**Supplementary Fig. 11**), whereas multiomic profiling enables definition of a refined, unified evolutionary trajectory by precisely delineating both cell identity and the boundary between normal and malignant states (**Fig. 4f**). DNA methylation changes are more conserved across individuals than transcriptomic alterations, as both PMD positions and the temporal order of PMD demethylation are largely consistent across patients (**Supplementary Fig. 10**; **Fig. 4e**). Together, these results highlight the value of multimodal approaches in capturing tumor evolution, with DNA methylation serving as a stable and generalizable molecular clock of disease progression.

### Intratumoral heterogeneity reveals a common origin of early PMD demethylation and somatic CNVs

We then asked, instead of building a pseudo-trajectory across samples, whether we could capture the evolution of the tumor genome and epigenome within the same tumor sample, to better reflect the true evolutionary trajectory. To this end, we quantified subclone-level hypomethylation fractions and identified two samples (RS35 and RS25 from G3) with heterogeneous methylation across subclones. Both samples comprised 3 aneuploid subclones, with one in each displaying a higher level of genome-wide methylation than the others (**Fig. 4g**).

To enable accurate inference of evolutionary relationships among subclones, we refined CNV calling by aggregating single-cell DNA reads into subclone-level pseudo-bulks, thereby enhancing the sensitivity for detecting CNVs shared across the majority of cells within each subclone. Following this refinement, we observed a clear separation between lineage-specific and shared CNVs (**Supplementary Fig. 12**), allowing confident reconstruction of phylogenetic trees that capture the clonal relationship among subpopulations within each patient sample.

In RS25, DNA methylation profiling revealed a progressive increase in hypomethylation across E1 and E2 PMDs among all the aneuploid subclones (**Fig. 4h**). Notably, although RS25_c3 and RS25_c2 exhibited methylation level comparable to those of euploid cells, this reduction was confined to a subset of E1/E2 PMDs. RS25_c2 displayed slightly greater PMD hypomethylation than RS25_c3, consistent with its more recent genetic divergence alongside the hypomethylated subclone RS25_c1. In RS35, a similar progressive increase in hypomethylation across E1 and E2 PMDs was observed, with more pronounced methylation loss in the progenitor-like subclones RS35_c2 and RS35_c3 (**Fig. 4h**).

To further clarify the temporal sequence between DNA hypomethylation and CNV alterations, we investigated high-versus low-methylation subgroups of luminal cells within each sample showing bimodality (**Supplementary Fig. 13a-b**). Cells with higher methylation largely exhibited intact diploid genomes, whereas a subset of hypomethylated cells remained CNV-negative, particularly in G2 relative to G3 samples (**Supplementary Fig. 13c**). We reasoned that this absence of CNVs may partially reflect limited sensitivity of CNV detection due to sparse single-cell DNA coverage, which would result in a higher false-negative rate in cells with lower CNV burden. Indeed, pseudo-bulk aggregation of hypomethylated LE cells from RS48 and RS32 revealed karyotypic changes on chromosome 6 and 8 (**Supplementary Fig. 14**), indicating coexistence of CNVs and global demethylation. However, RS7 from G2 showed no detectable CNV occurrence. The low fraction of hypomethylated cells, together with the absence of detectable CNVs in this sample, suggests a potential intermediate state in tumor progression, in which epigenetic disruption precedes large-scale genomic alterations.

These results suggest that canonical driver CNVs (e.g., 8p loss and 8q gain) arise in coordination with early DNA demethylation within PMDs. However, whether this coordination reflects a causal relationship or a consequence of selective pressure remains unclear and requires further investigation.

### Pre-neoplastic epigenetic burden as potential pioneer factor for tumor initiation

We next examined DMGs in epithelial cells to find out whether locus-level DNA methylation changes could be detected prior to CNV acquisition, aiming to identify potential pioneer events and epigenetic biomarkers of tumor initiation. Among the top 10 genes with the highest methylation gain, *TMEM106A* was consistently hypermethylated in G2 and G3 aneuploid LE cells relative to G1 coupled with downregulated expression (**Fig. 5a**), consistent with previous reports describing this gene as an epigenetic marker for PCa^28,29^. Intriguingly, promoter hypermethylation of *TMEM106A* was evident prior to the disease tipping point, with G3 cells showing a >20% increase in baseline methylation compared to G1 (**Fig. 5b**). The promoter of *HAAO*, another top 10 DMG, showed a similar but more modest baseline methylation, followed by a sharp increase coinciding with rising aneuploidy. Moreover, *TMEM106A* was found to be hypermethylated in G2 and G3 euploid cells relative to G1, albeit with a smaller effect size (<20%; **Fig. 5c**). These patterns suggested that *TMEM106A* hypermethylation may precede the acquisition of CNVs during tumorigenesis.

**Figure 5.**
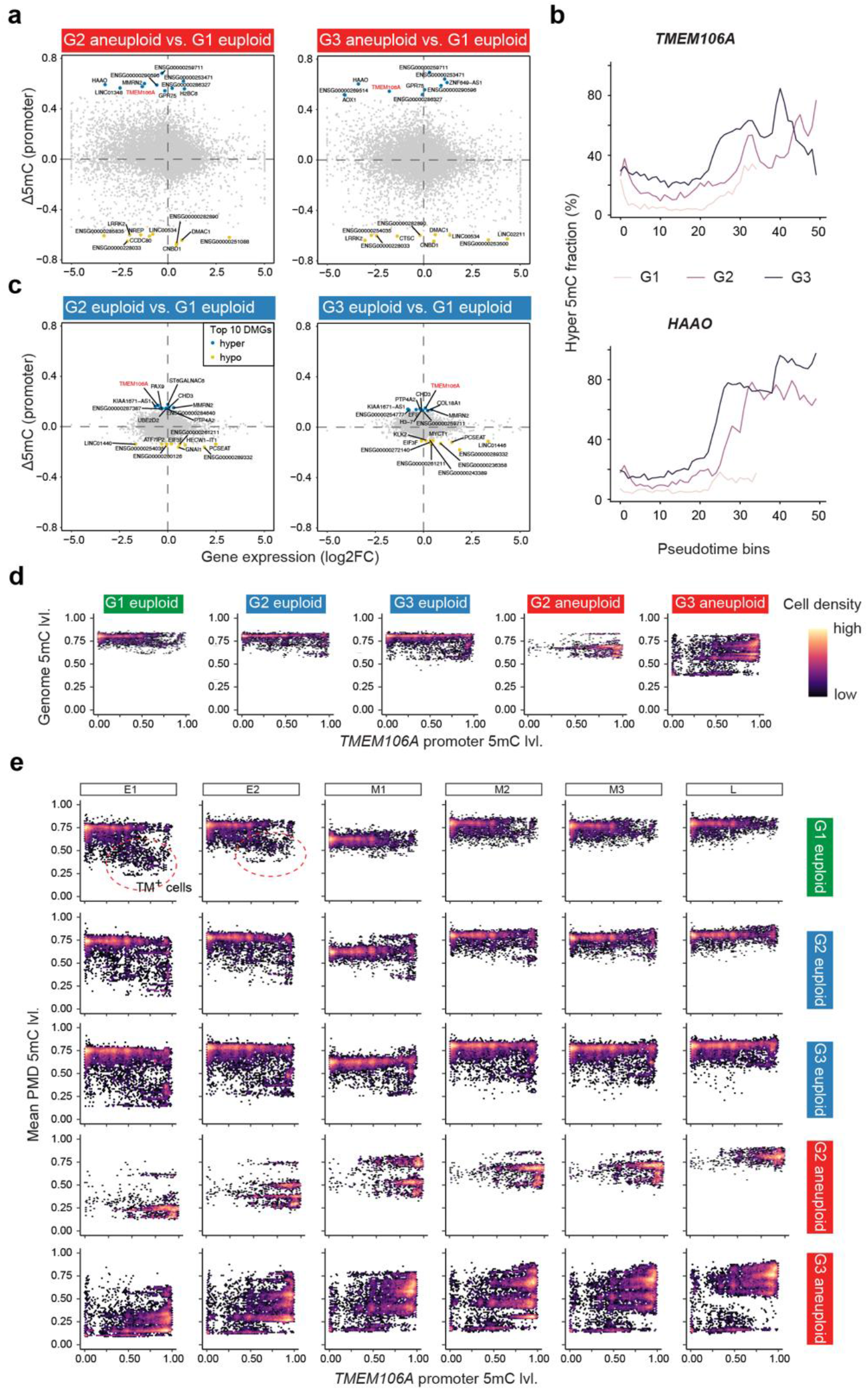
Tumor-associated DNA hyper- and hypomethylation patterns are evident in non-tumor BPH tissues. **a**, Changes in promoter methylation (Δ5mC) and gene expression (log_2_ fold change) in G2 and G3 aneuploid luminal epithelial (LE) cells relative to G1 euploid LE cells. **b**, Hypermethylated fraction (*y*-axis) for *TMEM106A* and *HAAO* promoter regions, defined as the proportion of cells with methylation level greater than 50%. **c**, Changes in promoter methylation (Δ5mC) and gene expression in G2 and G3 euploid LE cells relative to G1 euploid LE cells. **d**, Heatmap showing distribution of LE cells into different DNA methylation states (*x*-axis: *TMEM106A* gene promoter methylation; *y*-axis: the mean genome-wide 5mC level). **e**, Heatmap showing LE cells with different methylation patterns at cancer-related partially methylated domains (PMDs) and *TMEM106A* promoter. Red dashed circles highlight G1 TM^+^ cells that exhibited hypomethylated E1 and E2 PMDs and hypermethylated *TMEM106A*.

Given that both *TMEM106A* hypermethylation and early PMD hypomethylation are associated with the earliest CNV events in PCa samples (**Fig. 4h** and **Fig. 5a-c**), we asked whether their combination could serve as biomarkers of a pre-neoplastic cell state with epigenetic lesions. By jointly profiling *TMEM106A* and genome-wide methylation levels, we observed a clear progression from epigenomically intact cells (*TMEM106A* methylation low, genome methylation high) to hypomethylated states (*TMEM106A* methylation high, genome methylation low) (**Fig. 5d**, comparing G1 euploid against G2, G3 aneuploid).

Notably, a small fraction of hypomethylated cells was detected among G2 and G3 euploid populations (**Fig. 5d**). Based on this observation alone, we cannot exclude the possibility that these cells arise from false-negative CNV calls. However, we also identified hypomethylated cells in G1 samples (**Fig. 5e**, G1 euploid), which correspond to cancer-free prostatic tissues. These cells exhibited *TMEM106A* hypermethylation together with hypomethylation specifically at early PMDs (E1 and E2), hereafter referred to as TM^+^ cells, and did not yet display substantial loss of methylation at genome-wide level.

TM^+^ cells were found in 5 out of 15 benign samples (**Supplementary Fig. 15a**) and were preferentially found in patients with high-grade prostatic intraepithelial neoplasia (HGPIN; 2/2), despite the absence of histologically identifiable lesions in the scPRIME-seq profiled regions, compared to benign prostatic hyperplasia (BPH; 3/13) (**Supplementary Table 2**). We therefore propose that epigenetic degeneration may accumulate at variable rates in the normal prostate, and that early oncogenic events (e.g., 8p loss and 8q gain) preferentially arise in a cell context with pre-existing epigenetic alterations. This epigenetically primed state may facilitate the transition to HGPIN and subsequently to PCa.

Thus, through integration of single-cell multiomic evidence, we identified a rare pre-neoplastic epithelial state, termed TM^+^, characterized by tumor-permissive features and epigenetic alterations preceding the onset of multimodal collapse (i.e., the tipping point).

### Pre-neoplastic methylation changes are associated with altered transcriptomic state and microenvironmental interactions

To understand the functional consequences of pre-neoplastic methylation alterations and their potential link to tumorigenesis, we next focused on TM^+^ cells and examined their transcriptomic differences relative to normal epithelial cells, and interactions with other cell types within the pre-neoplastic microenvironment.

By comparing TM^+^ and TM^−^ cells, we found a largely consistent set of DEGs across G1, G2 and G3 euploid populations (**Fig. 6a**; **Supplementary Fig. 15b**). This coherent transcriptional pattern provides orthogonal validation for the methylation-based definition of TM^+^ cells, supporting their functional distinction relative to normal epithelium. GO analysis of the DEGs identified upregulation of synapse organization (GO:0050808), as well as downregulated antigen processing and presentation (GO:0019882) and stress response to metal ions (GO:0097501) (**Supplementary Table 5**). These changes are consistent with an immunosuppressive state and increased tolerance to oxidative stress and ferroptosis. The presence of these changes in G1 TM^+^ cells suggests the establishment of cellular state transition prior to malignant transformation.

**Figure 6.**
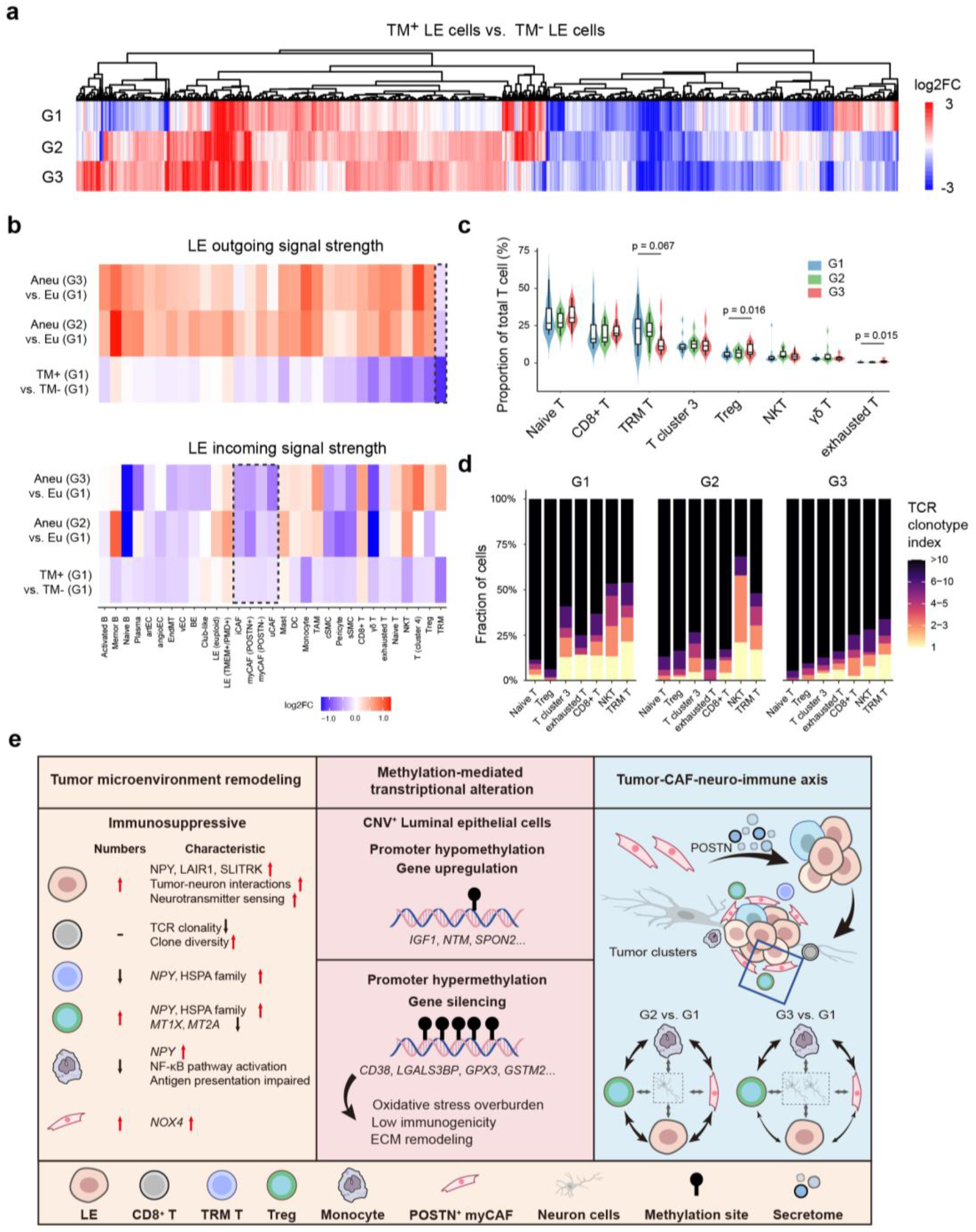
Cells with pre-neoplastic methylation changes adopt an altered transcriptional state and exhibit suppressive microenvironmental interactions. **a**, Differentially expressed genes between TM^+^ and TM^−^ cells are shown as a heatmap of log_2_ fold change. TM^+^ cells are defined by *TMEM106A* promoter methylation >50% and average E1/E2 PMD methylation <50%, with TM-cells defined conversely. Aneuploid LE cells were disregarded from G2 and G3. **b**, CellChat-inferred weighted signaling interactions between cell types. For each comparison (rows), log_2_ fold change in total interaction strength was computed between two populations across all partners. Dashed rectangles highlight consistent change in interactions of TM^+^ cells and cancer cells with TRM T cells (upper) and fibroblasts (bottom). **c**, Composition of T cell subclusters across patient groups. *P* values were calculated using two-sided Wilcoxon rank-sum tests between groups. **d**, TCR repertoire across G1-G3. Clonotype index represents the rank of TCR clonotype by their frequency within each patient sample. **e**, Summary of transcriptomic and epigenetic program changes during tumorigenesis, based on differential expression, gene ontology, CellChat and TCR repertoire analyses.

We next examined the cell-cell communication network of TM^+^ cells to assess their relationship with tumorigenesis. We compared TM^+^ and TM^−^ cells within G1, and contrasted this with communication patterns in G2/G3 aneuploid cells, using G1 euploid cells as a reference; the former captures pre-neoplastic TM^+^-specific alterations relative to normal epithelium, while the latter reflects broader tumor-associated microenvironmental changes. Although pre-neoplastic and tumor microenvironments are different, TM^+^ cells and aneuploid tumor cells consistently show a suppression of outgoing signaling to tissue-resident memory (TRM) T cells, together with reduced incoming signaling from fibroblasts (**Fig. 6b**), suggesting that loss of stromal regulation and immune interaction may already be established in the pre-neoplastic state. These results suggest that TM^+^ is a pre-neoplastic epigenetic state associated with tumor-permissive remodeling of intrinsic cell states that affects interactions with the microenvironment.

### Methylomic and transcriptomic states of advanced prostate cancer

Leveraging multiomic profiling by scPRIME-seq, we further assessed molecular alterations across the full spectrum of PCa from G1 to G3, by analyzing the 1,282/1,186 DEGs and 8,152/8,276 DMGs identified in G2/G3 aneuploid LE cells, respectively (**Fig. 2c**). Most DMGs reflected methylation loss, consistent with increased global hypomethylation from G2 to G3 (G2: 58.2%; G3: 74.5%), and were largely confined to heterochromatic regions, where they rarely led to gene activation (<1% genes upregulated; **Supplementary Fig. 8b**). In contrast, genes with detectable expression showed the expected inverse relationship between methylation and transcription, with >20% of DMGs overlapping inverse DEGs (**Supplementary Fig. 8c-d**), supporting a role for DNA methylation in modulating an active subset of the cancer transcriptome.

Integrative GO analysis further distinguished pathways with methylation-mediated regulation from those primarily reflecting transcriptional responses (**Supplementary Table 6**). Upregulated processes such as translation (GO:0002181) and oxidative phosphorylation (GO:0006119) showed minimal methylation changes (0-1 DMGs per term), whereas downregulated adhesion- and motility-related pathways (GO:0045785; GO:2000146; GO:0150115) in G2 showed stronger enrichment of hypermethylated genes (18.9-33.3% of DEGs; 5-8 DMGs per term), consistent with locus-specific methylation-associated repression. In G3, additional methylation-associated activation emerged in synaptic signaling (GO:0019883) and stress-response pathways (GO:0097501), while antigen presentation remained largely methylation-independent (GO:0019883). Overall, methylation-associated transcriptional effects were largely restricted to tumor epithelial cells (**Supplementary Table 6**).

In parallel, we profiled tumor microenvironment (TME) remodeling across PCa progression. G2/G3 tumors showed expansion of CNV^+^ LE cells and depletion of BE cells, alongside enrichment of POSTN^+^ myofibroblastic cancer-associated fibroblasts (myCAFs) and reduction of smooth muscle cells and monocytes (**Fig. 2c**). Cell-cell communication analysis revealed progressive strengthening of myCAF-derived signaling, whereas epithelial cells exhibited reduced incoming communication in G3, consistent with reduced stromal regulation (**Supplementary Fig. 16a**). Immune remodeling was characterized by depletion of TRM T cells and accumulation of Tregs and exhausted T cells (**Fig. 6c**), accompanied by reduced TCR clonality across CD8^+^, TRM and NKT compartments, consistent with an immunosuppressive shift (**Fig. 6d**).

Although neuronal populations were not captured in our dataset due to their incompatible cell size with droplet-based sequencing, additional network-level signals suggested activation of neuro-associated signaling programs, including NPY, SLITRK and SEMA pathways (**Supplementary Fig. 16b**). Correspondingly, G2 LE cancer cells exhibited hypomethylation and transcriptional activation of genes implicated in neuro-tumor interactions (*NTM, RAMP1, SPON2, and SYT14*) (**Supplementary Table 7**). G3 cancer cells displayed broader activation of neuro-associated programs, including genes involved in neuro-tumor interactions (*SPON2, LRRTM4, DMD*), neurotransmitter sensing (*OR51E2, CAMKK2*), and noncoding RNAs linked to neuro-tumor crosstalk (*FIRRE, CALCRL-AS*). Conversely, genes predicted to inhibit neuro-tumor interactions (*NDRG2, RASSF5*) were hypermethylated and downregulated. Also, *NPY* was upregulated in TRM, Treg, and monocytes; non-canonical NF-κB signaling was activated, including upregulation of *NFKB1*, *IL1B*, and *TNF* alongside downregulation of *MAPK14*, *JAK2*, and *MAP3K1* in monocytes showing reduced antigen-presentation genes (*CD36*, *IL15*), indicating a potential neurally influenced pro-inflammatory yet immunosuppressive state (**Supplementary Table 8**). POSTN⁺ myCAFs induce an oxidative stress microenvironment by upregulating *NOX4* expression. Additionally, the progressive enrichment of POSTN⁺ myCAFs from G1 to G3 (**Fig. 2c**) may further facilitate cancer cell clustering and perineural invasion^30,31^, as well as the formation of an immunosuppressive microenvironment (**Fig. 6e**).

Collectively, these results delineate an innervated and immunosuppressive tumor microenvironment in prostate cancer, where DNA methylation predominantly shapes cancer cell fate, while gene expression in stromal and immune cells is largely transcriptomically regulated.

## Discussion

The ability to read multiple modalities within the same cell is a fundamental requirement for understanding cellular processes, particularly in the context of cell-state transitions during disease initiation and progression. scPRIME-seq was developed to simultaneously profile genomic, epigenomic and transcriptomic information within individual cells, with substantially increased throughput (i.e., hundreds of thousands of cells) compared to existing methods. Through single-cell profiling of 44 human prostate samples, we reconstruct the evolutionary process from normal epithelium to malignancy, and provide a framework for multimodal biomarker discovery. Integration of CNV information not only enables more accurate delineation of tumor-non-tumor boundaries, but also provides additional resolution to map intratumoral heterogeneity, infer ancestral clones, and capture cellular trajectories along the benign-to-malignant transition. By resolving transitional cell states, our approach facilitates the identification of a “tipping point” during tumorigenesis, represented by a coordinated collapse across molecular modalities and irreversible shift from benign to malignant cell states (**Fig. 4f**).

Our results reveal that tumor cells accumulate epigenetic burden prior to the disease tipping point, as evidenced by PMD-associated hypomethylation and focal CpG island (CGI) hypermethylation (e.g., *TMEM106A* promoter) present in both tumor founder subclones and non-tumor cells (**Fig. 4h** and **Fig. 5e**). Mechanistically, DNA methylation loss has been linked to excessive cell division through passive dilution of 5mC^2^, and cancer-associated DNA hypomethylation is thought to arise primarily from passive rather than active demethylation, as indicated by the absence of 5hmC upregulation^32^. DNA methylation loss can occur in proliferative yet non-malignant contexts (e.g., BPH), or even in normal cell types without genomic alterations^2^, indicating that such systematic epigenetic alteration is not tumor-specific. This raises the possibility that PMD-associated hypomethylation reflects a pre-neoplastic epigenomic state, but is not sufficient on its own to drive malignant transformation. These findings challenge models in which epigenetic dysregulation arises primarily as a consequence of genomic instability during oncogenesis^33^, at least in prostate cancer.

It remains unclear what mechanisms drive the concurrent emergence of epigenomic and genomic alterations at the earliest stage of PCa. One possible explanation is that the normal epithelial cell state is inherently non-permissive to tumorigenesis, such that oncogenic potential of somatic mutations depends on prior epigenetic reprogramming to enable cell state transitions and microenvironmental shifts toward a tumor-permissive status. Indeed, early epigenetic alterations in PCa may already silence key tumor suppressor genes including *TMEM106A*^34,35^ and *CD1D*^7^, thereby preconditioning cells for subsequent oncogenic transformation. Supporting this hypothesis, a recent study demonstrated that chromosome 8 amplification promotes cellular fitness and oncogenesis in embryonic stem cells but not in differentiated neural progenitor cells^36^, suggesting that the oncogenic potential is indeed cell-state dependent. Alternatively, DNA methylation loss may directly lead to increased genomic instability through reactivation of transposable elements and increased DNA breaks^37,38^.

It is increasingly evident that DNA methylation alterations in cancer, especially DNA hypomethylation, may result in more complex functional consequences than immediate transcriptional changes. The observation that most hypomethylated regions remain transcriptionally inactive is consistent with a previous study identifying PMDs as being linked to widespread chromatin remodeling and deposition of gene-silencing-associated histone modifications (H3K27me3 and H3K9me3)^7^. We hypothesize that progression towards a low-methylation genomic state may provide increased epigenetic plasticity in tumors, in which gene silencing transitions from stable repression to more dynamic and potentially reversible states mediated by the Polycomb repressive complex (PRC). This increased plasticity may render tumor cells more adaptable to environmental stressors, including therapeutic pressures, and could facilitate state transitions associated with disease progression. Such a model may explain the emergence of alternative cellular states (e.g., neuroendocrine) during PCa therapy^39^.

Limitations remain. Although single-cell methylation sequencing enables the mapping of genomic reads and thus permits the inference of chromosome-arm-level somatic CNVs, accurate detection of single-nucleotide variants (SNVs) remains an obstacle due to low coverage and high dropout rates at the single-cell level. One potential solution is to couple scPRIME-seq with bulk whole-genome or whole-exome sequencing to generate a reference panel of somatic SNVs. Single-cell DNA reads from scPRIME-seq could then be mapped onto this panel to infer the distribution of SNVs at the subclonal level (e.g., by aggregating reads from each CNV subclone). In the context of PCa, however, we reason that our CNV-centric approach is unlikely to overlook key early oncogenic events, as tumor initiation is predominantly driven by CNV-related gene-level gains and losses^40^. Nonetheless, extending this strategy could be valuable for applying scPRIME-seq to tumor types in which tumorigenesis is primarily SNV-driven (e.g., gliomas^41^; melanoma^42^), and will require further methodological optimization and development of bioinformatics workflows.

In summary, scPRIME-seq provides a scalable workflow for characterizing cellular heterogeneity and dynamic transitional states, enabling the identification of intrinsic genetic and epigenetic determinants of transcriptomic programs and offering insights into the multimodal evolution of cancer.

## Methods

### Participants and biopsy procedures

We prospectively enrolled patients undergoing scheduled prostate biopsy. The inclusion criteria were as follows: (1) voluntary acceptance of prostate biopsy; (2) signing written informed consent approved by the institutional ethics committee; (3) agreement that harvested prostate specimens could be used for scientific research; (4) complete pre-biopsy serum PSA detection data; (5) intact preoperative imaging information; and (6) fully documented medical history and medication records.

Exclusion criteria included: (1) absolute contraindications to prostate biopsy, such as coagulation disorders and bleeding diathesis; (2) non-acinar adenocarcinoma pathological subtypes (e.g., small cell carcinoma); (3) active acute prostatitis; (4) history of prior prostate surgery; and (5) any other clinically inappropriate conditions judged by the investigators.

All biopsy specimens were acquired using an 18-gauge needle via transperineal MRI/TRUS fusion targeted biopsy combined with systematic prostate sampling. To ensure sufficient tissue specimens for routine clinical diagnosis and subsequent scPRIME-seq analysis, a parallel biopsy strategy was applied, with three tissue cores collected from identical focal lesions. Among them, one core was submitted for standard histopathological assessment including Gleason grading, while the remaining cores were reserved exclusively for translational research^43^. This study was ethically approved by the institutional review board (#2021-082) and strictly conducted in compliance with the principles outlined in the Declaration of Helsinki.

Participants were classified into three subgroups according to pathological diagnoses: G1 group with benign prostate histology, G2 group with low- to intermediate-risk prostate cancer group (acinar adenocarcinoma with Gleason score < 8, ISUP grade groups 1–3), G3 group with high-risk prostate cancer group (acinar adenocarcinoma with Gleason score ≥ 8, ISUP grade groups 4–5).

### Sample preparations

Tissue specimens were washed with pre-cooled RPMI-1640 medium and minced before being transferred into a centrifuge tube containing a digestion solution (1 mg/mL Collagenase II, 1 mg/mL Collagenase IV, 0.1 mg/mL DNase I, 1 mg/mL Dispase II, 2.5 mM CaCl_2_, and 2% fetal bovine serum). Digestion was performed in a hybridization oven at 37 °C for 30 min. Cell viability and concentration were assessed using acridine orange/propidium iodide (AO/PI) staining before the digestate was filtered through a 70 μm cell strainer. Cells were washed several times and finally resuspended in RPMI-1640 medium containing 2% FBS to achieve a concentration of 700–1,500 cells/µL.

### Cell fixation and permeabilization

Approximately 2×10^5^ cells were resuspended in 100 μL of PBS, and 400 μL of pre-chilled methanol was added dropwise to the cell suspension for fixation^25,44^. The mixture was subsequently incubated at –20 °C for 30 min before the cell pellet was sequentially washed with 1 mL of Wash Buffer A (3×SSC buffer [Sigma Aldrich, St. Louis, MO, USA; S6639], 1 U/μL RNase inhibitor, and 0.1% Tween-20) and Wash Buffer B (1× PBS, 1 U/μL RNase inhibitor, and 0.1% Tween-20).

### Intracellular chromatin tagmentation

Tagmentation was performed using a custom-assembled transposase, designated as Tn5 Enzyme A. The oligonucleotides used for the assembly of Tn5 Enzyme A (Yeasen, China, 12900ES60) were as follows: Tn5 oligo A: 5′-/5Phos/CTGTCTCTTATACACATCT-3′; Tn5 oligo B: 5′-/5Phos/CGTCCGTCGTTGCTCG/iBiodT/AGATGTGTATAAGAGACAG-3′; Tn5 oligo C: 5′-/5Biotin/GT/i5MedC/T/i5MedC/GTGGG/i5MedC/T/i5MedC/GGAGATGTGTATA AGAGA/i5MedC/AG-3′.

Approximately 3×10^4^ cells were transferred into 60 μL of reaction buffer containing 11 U of Tn5 Enzyme A. The mixture was incubated at 37 °C for 60 min. Following incubation, the samples were washed with PBS containing 0.1% Tween-20 and resuspended in 50 μL of PBS supplemented with 1 U/μL RNase inhibitor.

### Single-cell nucleic acid barcoding inside droplets

The barcoded sequence immobilized on the gel beads was designed with the following structure: AmCGAmCGmCTmCTTmCmCGATmCT(j)_17_(D)_12_TTTCTTATATrGrGrG, where (j)_17_ represents the cytosine-free cell barcode, and (D)_12_ represents the unique molecular identifier (UMI) sequence, which is also cytosine-free. The terminal segment, TTTCTTATATrGrGrG, functions as a switch oligonucleotide to facilitate DNA conjugation. For mRNA capture, the reverse transcriptase incorporates the gel bead oligonucleotide at the 5′ end of the transcript via a template-switching reaction.

Cells were mixed with a ligation and reverse transcription (RT) system. The reaction mixture contained: 8 μL of 10× Ligation Buffer (Enzymatics, MA, US; B6030L); 1 U RNase inhibitor; 1 mM dATP/dTTP/dGTP; 1 mM 5-methyl-dCTP (New England Biolabs, MA, US; N0356S); 15 mM DTT; 3.2 μL of T4 RNA Ligase 2 (New England Biolabs, MA, US; M0239S); 3.2 μL of RT Enzyme (SeekGene Biosciences, China; R0003802); RT primer (5′-AAGCAGTGGTATCAACGCAGAGTACTTTTTTTTTTTTTTTTTTTTTTTTTTTTTTVN-3′); 0.3 mM bridge sequence (5′-AGCAACGACGGACGCCCATATAAGAAA-3′). Droplets were generated using the SeekOne® Digital Droplet Instrument (SeekGene Biosciences, China; M001A). The emulsion was transferred to a new PCR tube and incubated at 25 °C for 30 min, followed by 42 °C for 60 min to perform concurrent ligation and reverse transcription. To terminate the reaction, 3 μL of 10% SDS was added to the emulsion. The oil phase was carefully removed, and the aqueous phase containing the nucleic acids was purified using clean-up beads (SeekGene Biosciences, China; R0003402).

### Separation of gDNA and cDNA

The separation of genomic DNA (gDNA) and cDNA was performed using Dynabeads™ M-280 Streptavidin magnetic beads (Thermo Fisher Scientific, MA, US; 11205D) according to the manufacturer’s protocol. The DNA bound to the magnetic beads was retained for subsequent DNA methylation library preparation, while the cDNA remaining in the supernatant was collected and purified using VAHTS DNA Clean Beads (Vazyme Biotech, China; N411-03) for 5′ transcriptome library construction.

### Single-cell DNA methylation library construction

The transposed DNA was first subjected to a gap-filling reaction by adding 50 µL of a gap-fill mixture and incubated at 65 °C for 20 min. To prevent the conversion of the cell barcode and functional sequences (e.g., sequencing-primer regions) during the subsequent enzyme-mediated conversion of cytosine bases, the gap-fill reaction mixture exclusively utilized 5-methyl-dCTP and omitted dCTP.

The DNA was then oxidized with TET2 and deaminated with APOBEC using the NEBNext® Enzymatic Methyl-seq Kit (New England Biolabs, MA, US; E7120S) according to the manufacturer’s instructions. Following C-to-U conversion, the DNA was amplified using the following primers (asterisks denote phosphorothioate bonds): 5′-CTACACGACGCTCTTCCGATCT-3′ and 5′-GTCTCGTGGGCTCGGA*G-3′. The resulting PCR product was purified using 0.8× VAHTS DNA Clean Beads and further fragmented using a Tn5 transposome. This Tn5 transposome was pre-assembled by loading the Tn5 transposase (Tn5 enzyme B) with annealed adapter oligonucleotides: 5′-GTCTCGTGGGCTCGGAGATGTGTATAAGAGACAG-3′ and 5′-/5Phos/CTGTCTCTTATACACATCT-3′. For fragmentation, 100 ng of purified DNA was incubated with 1 U of the pre-assembled Tn5 transposome at 55°C for 10 minutes. The purified library was amplified with TruSeq P5 and Nextera P7 index primers.

### Single-cell 5′ transcriptome library construction

To minimize non-target DNA amplification, cDNA purification was conducted using a two-round PCR strategy.

In the first round, purified cDNA was amplified with the primers 5′-CTACACGACGCTCTTCCGATCT-3′ and 5′-AAGCAGTGGTATCAACGCAGAGT-3′ using the following program: 98 °C for 3 min; 7 cycles of 98 °C for 30 s, 63 °C for 15 s, and 72 °C for 2 min; final extension at 72 °C for 5 min, and hold at 4 °C. The product was purified using 0.8×VAHTS DNA Clean Beads.

A second PCR was then performed using an aliquot of the purified cDNA, maintaining the exact reaction components and cycling parameters as the first round. The resulting cDNA was purified with 0.6×VAHTS DNA Clean Beads and fragmented using the Nextera Library Preparation Kit under the identical conditions described for the DNA library. Following fragmentation, library amplification was carried out as described for the DNA methylation library. The final 5′ transcriptome library was purified using 0.8×VAHTS DNA Clean Beads.

For T-cell receptor (TCR) profiling, libraries were prepared utilizing the SeekOne™ DD Single-Cell TCR Enrichment Kit (Human) (SeekGene Biosciences, China; K00601) according to the manufacturer’s protocol. Briefly, the second-round amplified cDNA underwent two successive enrichment PCRs using human TCR primers 1 and 2. Following fragmentation and adapter ligation, the TCR library was amplified with TruSeq P5 and P7 index primers (98 °C for 3 min; 8 cycles of 98 °C for 20 s, 54 °C for 30 s, and 72 °C for 20 s; final extension at 72 °C for 5 min, hold at 4 °C). The final TCR library was purified with 0.8×VAHTS DNA Clean Beads.

### Sequencing

The purified libraries were sequenced on the SURFSeq 5000 platform (GeneMind Biosciences LTD, China) utilizing a 150-cycle paired-end, high-output sequencing strategy.

### RNA and DNA methylation sequencing data analysis

The paired single-cell RNA-methylation sequencing data were processed using the SeekSoulMethyl pipeline developed by SeekGene (v1.0.6; https://github.com/seekgene/SeekSoulMethyl) to generate the transcriptomic expression matrix and the single-cell methylation matrix.

First, raw FASTQ files from the transcriptomic and methylation libraries were subjected to quality control separately. The transcriptomic libraries were processed using fastp^45^ (v1.0.1) for basic quality control and trimming of low-quality bases. The methylation libraries were also preprocessed using fastp, with automatic adapter trimming disabled in order to preserve the fixed library structure required for subsequent cell barcode and UMI identification.

The transcriptomic branch was processed using SeekSoul Tools (v1.3.0; https://github.com/seekgenebio/seeksoultools) for sequence alignment, gene quantification, and expression matrix generation. Internally, reads were aligned using STAR^46^ (v2.7.10a), and a cell-filtered expression matrix was generated. Intronic reads were retained during quantification to improve the sensitivity of single-cell RNA detection. For different library chemistries, the pipeline automatically matched the corresponding RNA analysis parameters: DD-MET5 samples were processed with DD-MET5, whereas DD-MET3 samples used DDV2 for the transcriptomic branch.

For the methylation branch, cell barcodes and UMIs were first identified and corrected according to the library structure, and the corresponding barcode whitelist was selected based on the chemistry type (DD-MET3 or DD-MET5). If an extracted barcode was not present in the whitelist but differed from a whitelist entry by a Hamming distance of 1, barcode correction was attempted; reads with uncorrectable barcodes were discarded. In the same step, read orientation was determined based on the fixed library sequences, the C-to-T conversion rate was estimated, and artificial sequences introduced during library construction were removed, including the TSO/7F linker, 17L and ME sequences, as well as the 9-bp gap introduced by Tn5 transposition, in order to reduce artificial methylation signals in downstream analyses. Read pairs with excessive non-CpG methylated cytosines were filtered out; specifically, a read pair was removed when more than two non-CpG methylated C bases were detected. In addition, only reads with a processed Read1 length of at least 20 bp and a processed Read2 length of at least 60 bp were retained for downstream analysis.

The preprocessed methylation reads were aligned using Bismark^47^ (v0.25.2). In this workflow, we modified Bismark by introducing the --add_barcode and --add_umi options, which write CB (corrected cell barcode) and UR (raw UMI) tags, respectively, into the BAM file to support downstream per-cell splitting and UMI deduplication. Forward reads were aligned with a maximum insert size of 1000 bp, whereas reverse reads were aligned in PBAT mode using the --pbat and -X 1000 parameters. After alignment, BAM files were sorted by read name using samtools to prepare for subsequent single-cell splitting.

During single-cell methylation matrix construction, methylation BAM files were first split by cell according to the list of high-confidence cell barcodes obtained from the transcriptomic analysis, and only methylation cells matching the high-confidence transcriptomic cell barcodes were retained. Forward and reverse BAM files corresponding to the same cell were then merged to generate single-cell BAM files, which were subsequently converted to ALLC files using ALLCools^21^ (v1.2.0). In this step, ALLCools performed UMI correction based on the UR tag and deduplication at the individual cytosine level. ALLCools was then used to construct multi-scale single-cell methylation matrices, including matrices based on multiple genomic binning scales and gene-neighborhood scales. The final single-cell methylation matrix was output in .mcds format.

### Clustering analysis and cell type annotation

Both transcriptomic data and DNA methylation data were used to compute principal component embeddings and identify cell clusters using the Leiden algorithm, enabling comparison between modalities. All downstream analyses were based on transcriptome-derived clustering for cell type annotation.

For transcriptomic data, cells were filtered to retain those with 200-5,000 detected genes and less than 20% mitochondrial reads. Raw counts were normalized, log-transformed, and scaled, and the top 4,000 highly variable genes were used for principal component analysis (PCA). Data integration across samples was performed using the Python implementation of Harmony with ‘nclust’ set to 50.

Cell clusters were identified using a consensus clustering strategy in the harmonized PCA space. Briefly, a baseline Leiden clustering was first performed to assign initial cluster labels. This was followed by 20 bootstrap Leiden clustering runs with different random seeds, and cluster labels were aligned to the baseline. Final cluster assignments were determined by majority voting across bootstrap iterations.

For methylation data, clustering was performed using a binarized 20-kb genomic bin matrix without additional imputation, where missing values were assigned by ALLCools^21^ based on cell-level mean methylation.

Cell types were annotated based on expression of canonical marker genes. Secondary clustering (e.g. subclusters within epithelial and immune cells) was performed using the same consensus clustering framework.

### Methylation data missing value imputation via PRIME-IMPUTE

Imputation of missing values was performed starting from the .mcds output generated by ALLCools^21^. Two distinct strategies were applied depending on genomic feature type.

For gene-associated regions, including gene bodies with ±2 kb flanking regions (geneslop2k), and promoter regions, raw methylation levels were defined as the fraction of methylated CpG sites within each region (ranging from 0 to 1). Missing values in these regions were initially assigned by ALLCools as the average methylation fraction across CpG sites within the same cell.

For the genome-wide 20 kb bins (chrom20k), raw methylation values were first normalized into the relative ratio with respect to the cell-specific mean methylation level. Missing values were initialized to 1, indicating no deviation from the cell mean. These normalized values were subsequently binarized into a 0-1 representation, where 0 denotes the methylated state and 1 denotes the unmethylated state, based on whether the value falls below or above the cell mean. Following these preprocessing steps, all missing values were explicitly relabeled as −1.

Imputation was conducted at the per-sample level, such that only cells derived from the same sample were used to infer missing values. A k-nearest neighbors (kNN) graph was constructed using the top 20 principal components (PCs) derived from PCA embeddings of the gene expression matrix across all cells.

For a given missing value *X* in a target cell, imputation was performed by estimating a weighted average of the corresponding feature values from its *k* nearest neighboring cells:

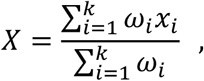

where *x_i_* denotes the methylation values of the *i* th neighboring cell, and *ω_i_* represents its corresponding weight. Weights were defined using an inverse distance:

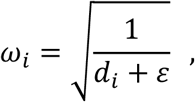

where *d_i_* is the Euclidean distance the target cell and its neighbor *i* in PCA space, and *ε* is a small constant introduced to avoid numeric instability.

For imputing binarized methylation states (chrom20k), the weights of neighboring cells were aggregated separately for each class (0: methylated; 1: unmethylated). The imputed label for each missing entry was then assigned to the class with the higher total weight.

Only neighbors with valid non-missing methylation values were considered, and imputation was performed only if at least *m* of the *k* nearest neighboring cells contained non-missing values for that feature. The parameters *k* and *m* were adaptively determined according to feature type (promoter, geneslop2k, or chrom20k).

### Differential gene expression analysis

Differentially expressed genes (DEGs) were identified using the Python Scanpy package^48^, applying a Wilcoxon rank-sum test for both one-vs-rest and pairwise comparisons between cell groups. DEGs were defined based on the following criteria: (1) Benjamini-Hochberg-adjusted *p*-value < 0.05; (2) absolute log_2_ fold change > 1; and (3) expression detected in at least 10% of cells in either the target or the reference cell group.

### Differential methylation analysis

Differentially methylated genes (DMGs) or regions (DMRs) were identified across multiple genomic feature levels (promoter, geneslop2k, chrom20k). For continuous methylation values, statistical significance was assessed using a Wilcoxon rank-sum test, whereas binarized methylation states were evaluated using Fisher’s exact test. For each comparison, the difference in methylation level (Δ5mC) was calculated as the absolute difference in mean methylation between the groups. DMGs/DMRs were defined as features with an adjusted *p*-value < 0.05 and an |Δ5mC| > 0.15 (i.e., a minimum 15% difference in methylation levels), after excluding sites with <10% methylation calling rate (e.g., low coverage) in either the target or reference cell group.

### Copy number variation calling

CNV analysis was performed based on single-cell methylation BAM files generated after per-cell splitting and merging. First, the FLAG information for Read1/Read2 in reverse-orientation BAM files was corrected to standardize downstream parsing of paired-end reads. The BAM files were then sorted and indexed using SAMtools^49^, followed by UMI deduplication using UMI-tools^50^ (v1.1.6) based on the UR tag. In this step, UMIs were extracted from BAM tags and deduplicated in paired-end mode; the parameters --umi-tag UR, --paired, and --ignore-tlen were used to accommodate the tag structure and paired-read characteristics of the present dataset. The deduplicated single-cell BAM files were used for downstream CNV inference.

CNV calling was subsequently performed using CopyKit^51^ (v0.1.3). The data were treated as paired-end data, and remove_Y = TRUE was applied to reduce the influence of chromosome Y signals on CNV inference. CNV analysis was carried out at the binning resolution of 1 Mb with the default parameter min_bincount = 10. The resulting heatmaps of subclonal CNV distributions were manually inspected, and low-confidence subclones, defined as those supported by singleton CNV events and lacking intrachromosomal breakpoints (likely arising from read dropout artifacts), were excluded from the aneuploid subclone list.

### Copy number variation pseudo-bulk correction

For a given list of cell barcodes presumed to originate from a homogeneous population (e.g., cells within a CNV-defined subclone), a refined pseudo-bulk analysis was performed to assess potential false-positive and false-negative calls from single-cell level CNV inference. Genomic reads associated with the selected cell barcodes were extracted from individual BAM files and merged into a single BAM file, representing a collective pseudo-bulk sample. Per-base read coverage of the pseudo-bulk sample was then calculated using “samtools depth -a” and subsequently aggregated into 250-kb genomic bins across chromosomes. As a control, an equal number of randomly selected cell barcodes lacking CNVs were extracted from the same source (i.e. same RS ID) to construct a diploid reference. For each 250-kb window, the ratio of coverage between the test and control samples, multiplied by two, was used to estimate regional ploidy (with the exception of chromosome X, representing the only haploid chromosome in the male genome). The resulting genome-wide coverage distribution was manually inspected to validate pseudo-bulk-level aneuploidy patterns.

### TCR and BCR analysis

The raw sequencing data were processed by fastp^45^ to trim primer sequences and low-quality bases. The remaining data was demultiplexed based on cellular barcodes and mapped reads to the human GRCh38 reference genome using SeekSoul Tools (Seekgene, v1.3.0) pipeline to generate a feature-barcode matrix, which was subsequently converted into a Seurat object in R used for the downstream analyses.

For V(D)J libraries, Seeksoul tool (Seekgene, v1.3.0) was used to perform demultiplexing, *de novo* assembly, contig annotations, clonotype calling, and V(D)J read alignment to the manufacturer-supplied human V(D)J reference genome.

### Partially methylated domain (PMD) identification

PMD identification was performed using binarized, missing-value-imputed methylation data matrix at 20-kb genomic resolution (chrom20k). PMD analysis was restricted to LE cells. The procedure comprised sample-level PMD calling followed by cross-sample consolidation of shared PMDs.

At the sample level, genome-wide methylation levels were scanned using a sliding window of 100 kb (5 consecutive 20-kb bins) with a step size of 20 kb. For each window, a hypomethylation fraction was calculated as the average fraction of cells exhibiting a hypomethylated state (binary encoding) across constituent bins, excluding missing values. Windows with a hypomethylation fraction ≥0.6 were flagged as hypomethylated. Overlapping windows were merged to define candidate PMDs, and regions spanning fewer than 13 bins (<260 kb) were excluded.

To derive a unified set of PMDs across samples, domains identified independently in each sample were consolidated based on a minimal 70% reciprocal overlap. A subsequent iterative refinement step was applied to merge smaller PMDs into larger PMDs that covered at least 80% of their length. PMDs not overlapping others were retained as singleton domains. The resulting 1,338 PMDs were used for downstream analysis.

### Gene ontology (GO) analysis

Genes meeting the DEG criteria (|log_2_ fold change| > 1 & false-discovery rate < 0.05) in pairwise G1/G2/G3 comparisons for each cell type were used for GO analysis. Upregulated and downregulated genes were analyzed separately using the “enrichGO” function from the ‘clusterProfiler’ R package, with the Benjamini-Hochberg (BH) method for multiple testing correction and a adjusted *p* value cutoff of 0.05. For significantly enriched GO terms, we quantified the overlap between DEGs assigned to each term and DMGs, considering only DMGs with inverse directional changes relative to the corresponding DEGs. The fraction of DEGs overlapping DMGs was calculated to estimate the contribution of epigenetic remodeling to the observed alterations in biological processes.

### CellChat analysis

CellChat^52^ (v2.1.2) was used for the analysis. Cell-cell interaction networks were first inferred separately for G1, G2 and G3, and subsequently merged for pairwise comparisons.

## Acknowledgements

We thank Prof. Jinsong Li (Shanghai Institute of Biochemistry and Cell Biology, CAS) for helpful scientific discussions, and thank the staff member Pengyu Wang of Bio-Med Big Data Center/Shanghai Institute of Nutrition and Health (SINH) for providing invaluable technical support and assistance in data collection.

## Author contributions

Conceptualization: Z.F.L., S.H., and S.L.

Methodology: S.L., Y.X., X.M., and M.T.

Investigation: S.L., Z.Q.L., X.W., H.Y., S.H., and D.W.

Visualization: S.L. and Z.Q.L.

Funding acquisition: Z.F.L., S.H., and D.W.

Supervision: Z.F.L., S.L., and S.H.

Writing: S.L., Z.F.L., and Z.Q.L.

## Supplementary figures

**Supplementary Figure 1.**
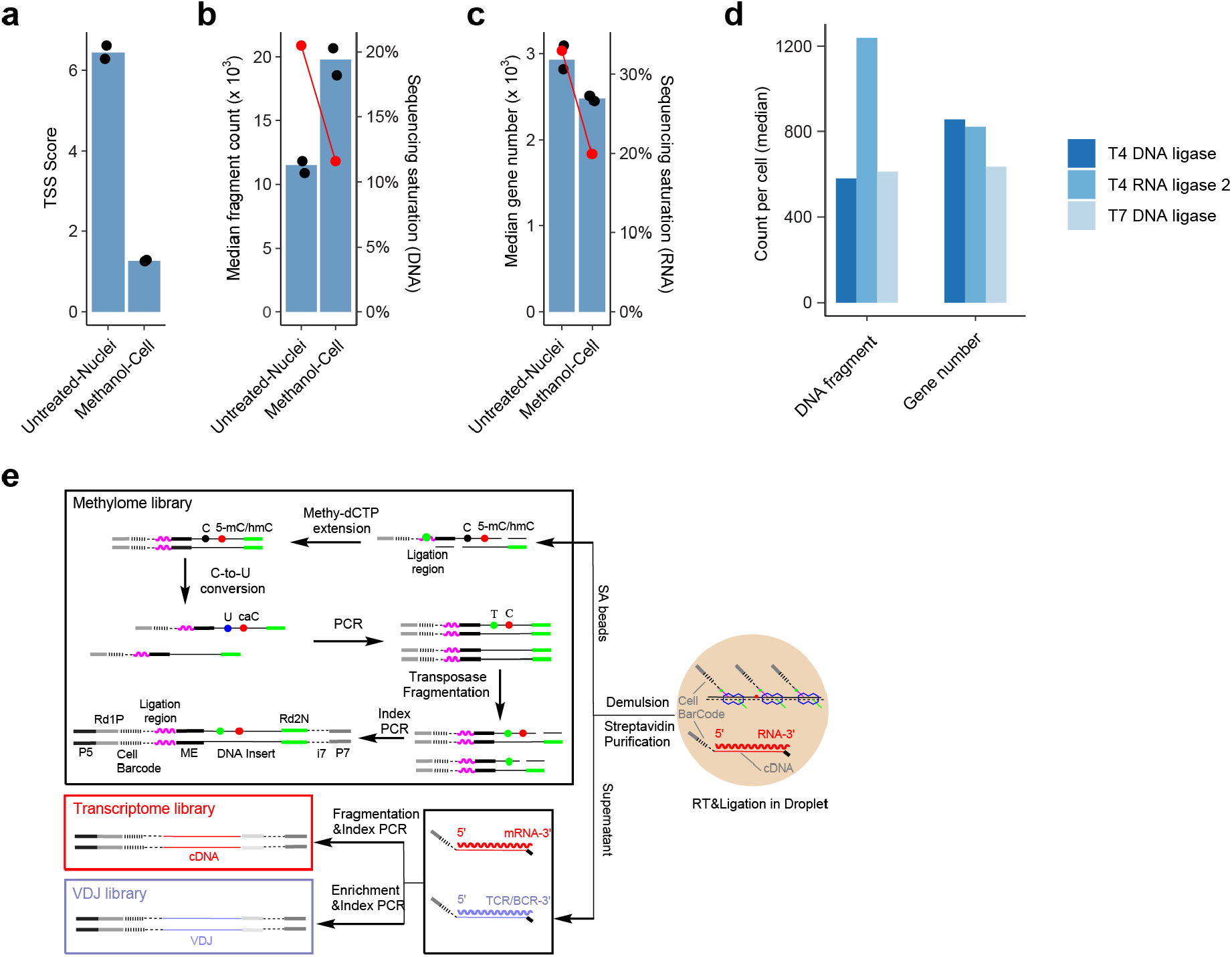
Optimization of scPRIME-seq workflow. **a**, Transcription start site (TSS) score calculated for whole-cell samples treated with methanol, compared to extracted nuclei without treatment. A TSS score of 1 denotes even distribution of reads between TSS region and other genomic regions, whereas a positive TSS score indicates coverage bias towards TSS (i.e. open chromatin). **b**, Median number of DNA fragments captured per cell. Red dotted line indicates the sequencing saturation rate for each condition. **c**, Median number of genes with detected transcription per cell. **d**, Comparison of DNA and RNA ligases for performance in methylome library construction. **e**, Schematic illustration of library preparation steps following demulsion of the droplets.

**Supplementary Figure 2.**
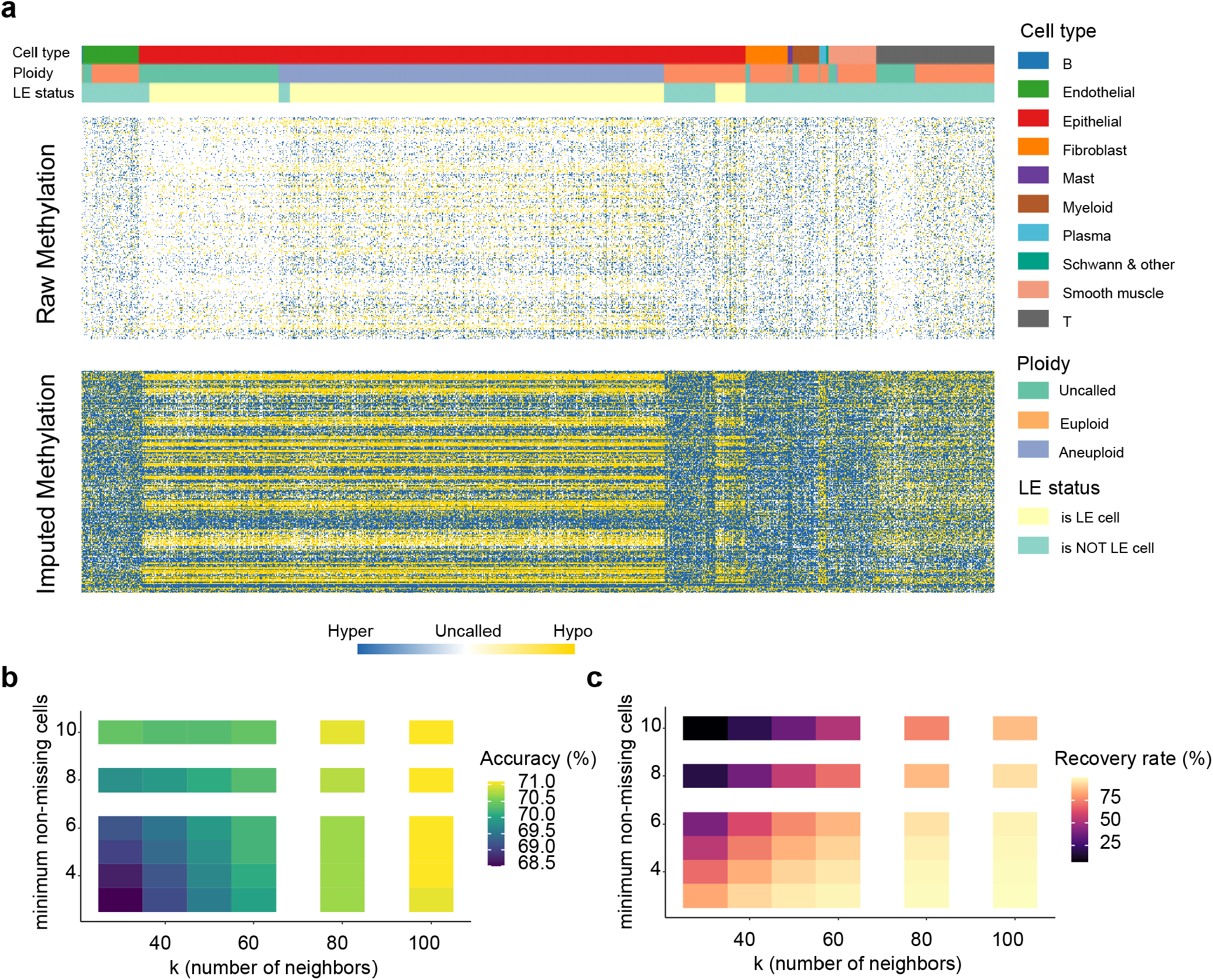
Transcriptome-guided imputation of sparse single-cell methylome data. **a**, Application of transcriptome-guided DNA methylation imputation to a patient dataset (RS67). Heatmap showing binarized methylation states for 20kb genomic bins across chromosome 8 (rows) in all cells (column) from RS67. **b**, Grid search for optimal imputation hyperparameters. Accuracy was calculated by masking 10% of observed methylation values and re-imputation. **c**, Recovery rate of missing methylation values across different hyperparameter combinations.

**Supplementary Figure 3.**
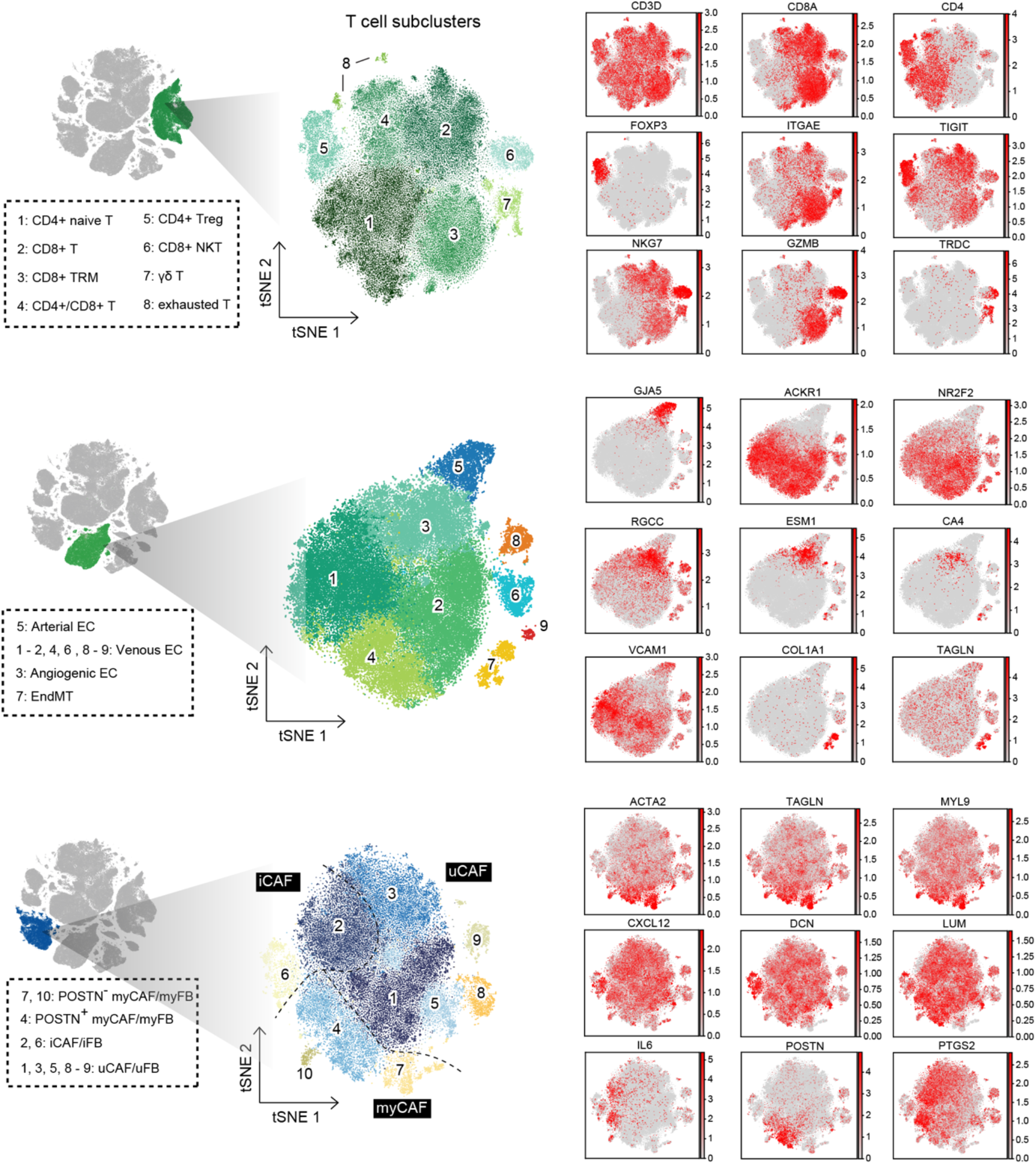
Subcluster annotation of T cells, endothelial cells and fibroblasts. Secondary consensus clustering was performed for T cells, endothelial cells and fibroblasts. Subclusters were then annotated based on marker gene expression (right panels for each cell type). In the gene expression panels, color intensity denotes expression level, with grey indicating no detectable expression.

**Supplementary Figure 4.**
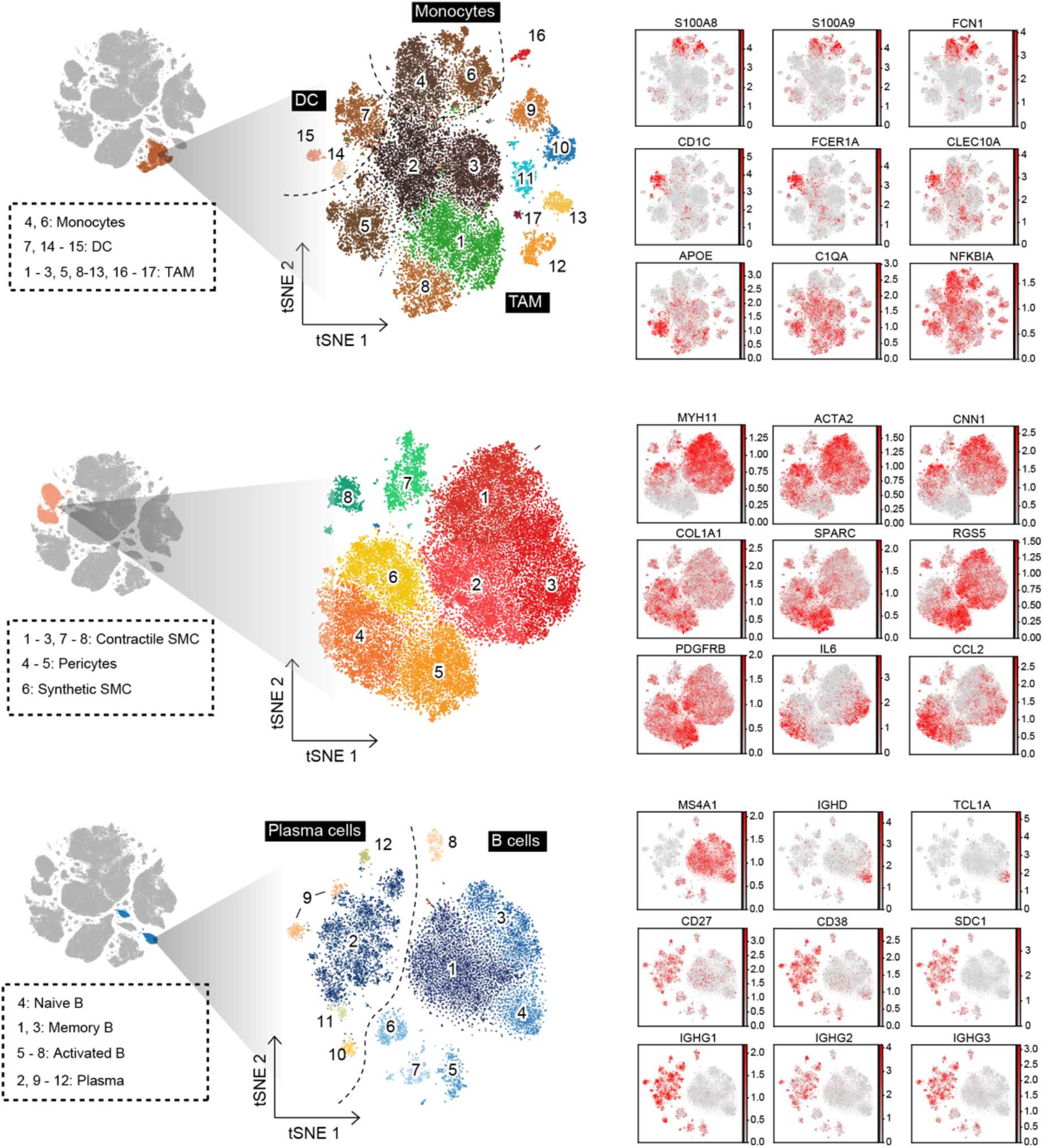
Subcluster annotation of myeloid cells, smooth muscle cells and B cells. Secondary consensus clustering was performed for myeloid cells, smooth muscle cells (SMCs) and B cells. Subclusters were then annotated based on marker gene expression (right panels for each cell type). In the gene expression panels, color intensity denotes expression level, with grey indicating no detectable expression.

**Supplementary Figure 5.**
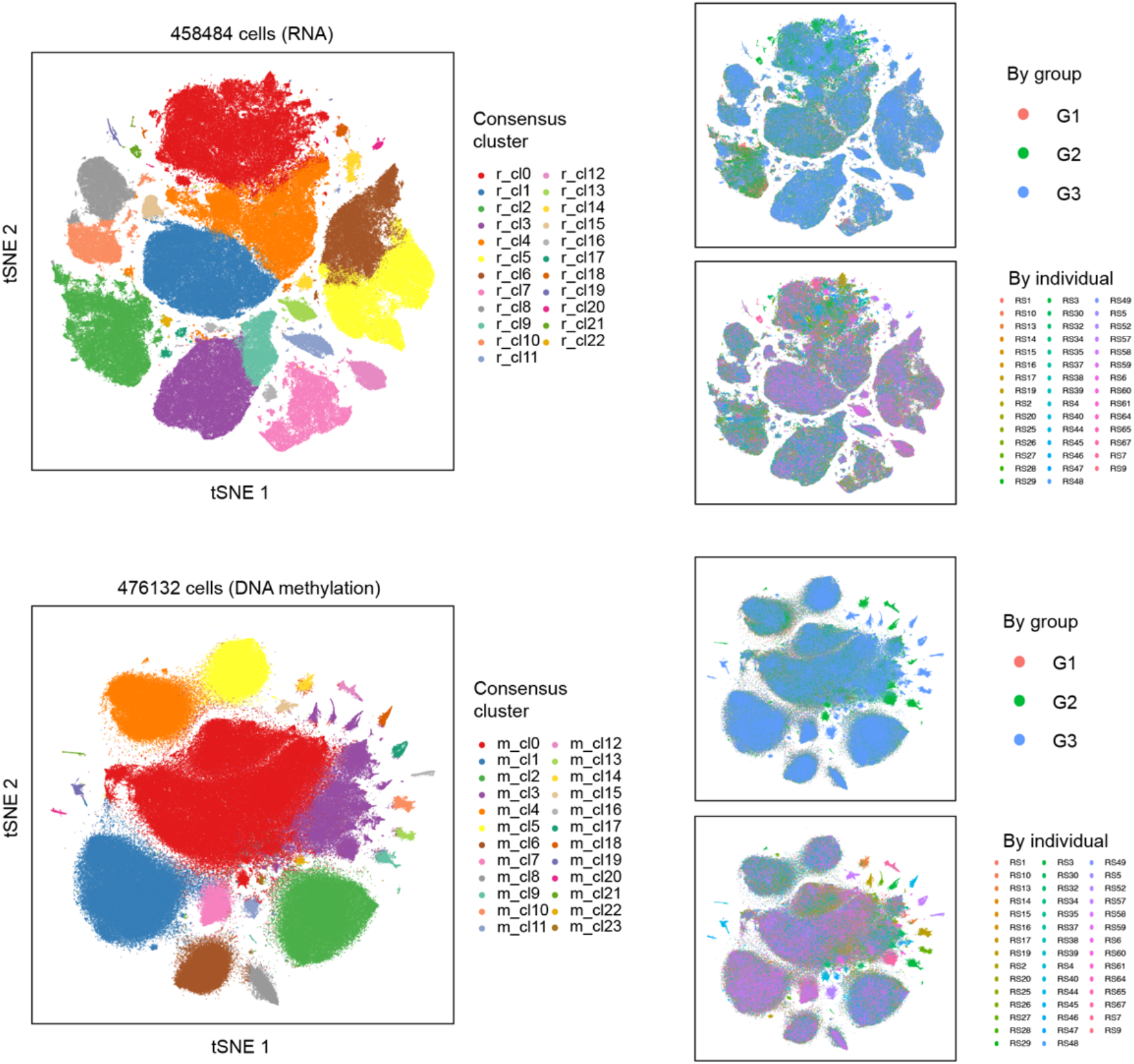
Transcriptome- and methylome-based Leiden clustering. t-SNE embeddings generated independently from transcriptome (upper) and methylome (lower) data for all valid cells across 44 patient samples. Methylome clustering was performed using chrom20k data without transcriptome-guided imputation.

**Supplementary Figure 6.**
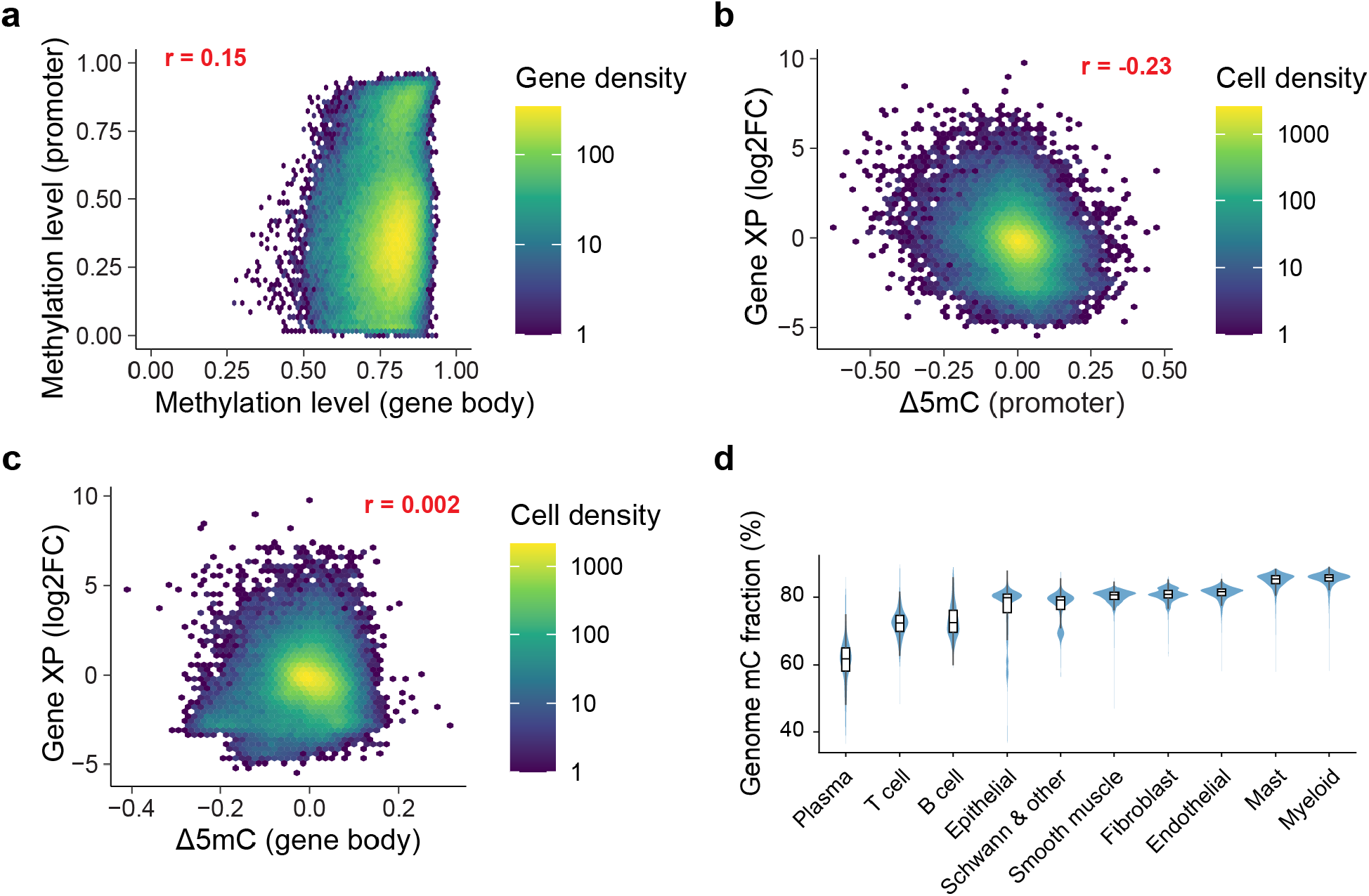
Relationship between DNA methylation and transcription at single-cell resolution. **a**, Distribution of methylation levels in gene bodies (geneslop2k; TSS to TTS ± 2 kb) and promoters; *r* denotes Pearson’s correlation coefficient. **b-c**, Distribution of Δ5mC from 1-to-rest DMG analysis across 10 major cell types, compared with corresponding gene expression changes. Panel **b** shows promoter Δ5mC and panel **c** shows gene body Δ5mC. Only genes with detectable expression in at least one cell type and with length > 15kb were included (to avoid confounding from flanking regions in gene body methylation estimates using geneslop2k). **d**, Distribution of genome methylation level across 10 major cell types.

**Supplementary Figure 7.**
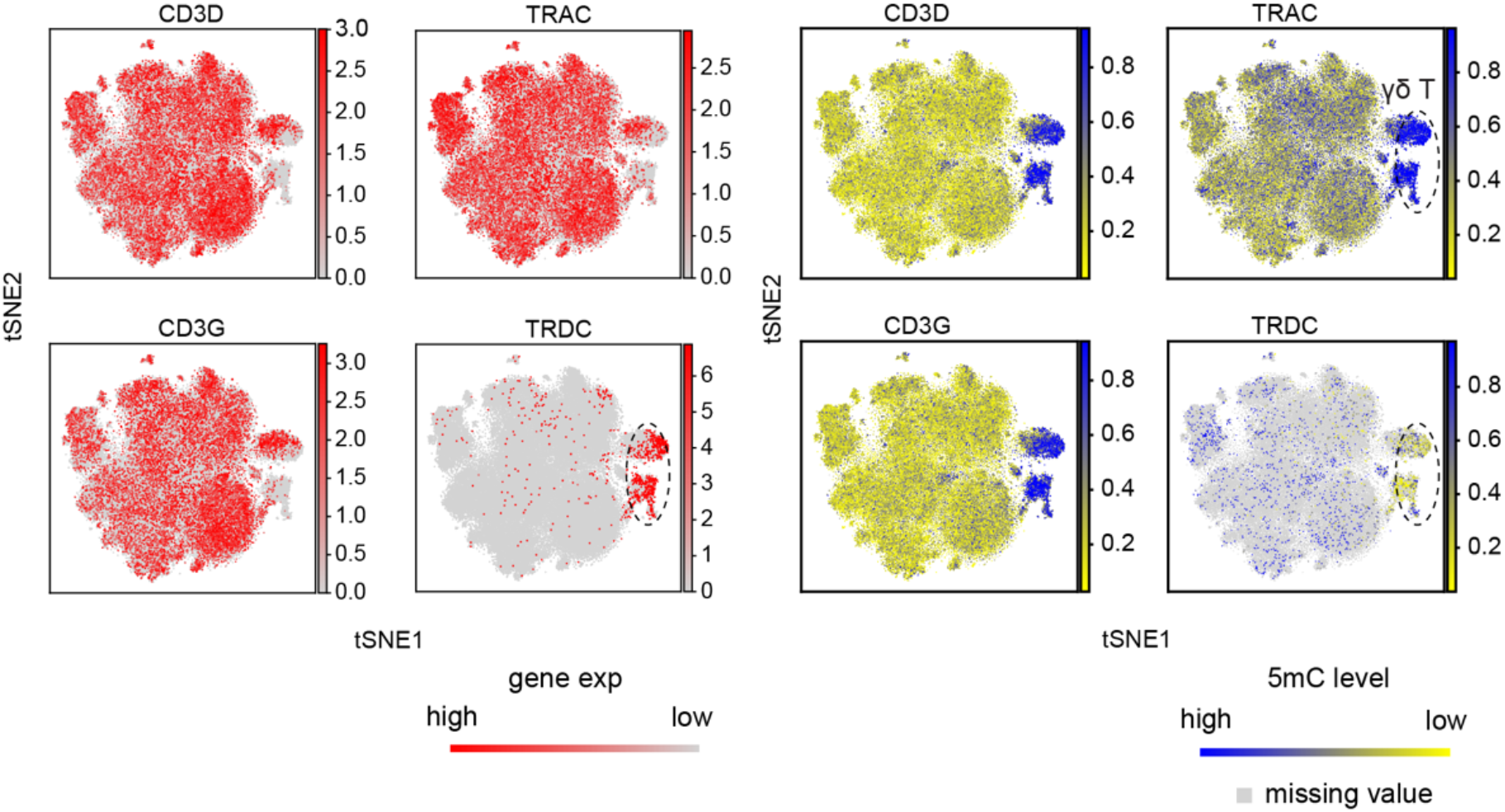
Identification of a CD3^−^ γδ T cell cluster using single-cell transcriptome and methylome evidence. Gene expression (left) and promoter region DNA methylation (right) distributions are overlaid onto the t-SNE embedding space of all T cells across 44 samples. For the methylation data, cells with missing values (after PRIME-IMPUTE imputation) are indicated in grey, representing either insufficient coverage or true depletion of DNA fragments due to TCR locus rearrangement (e.g. *TRDC* deletion in αβ T cells).

**Supplementary Figure 8.**
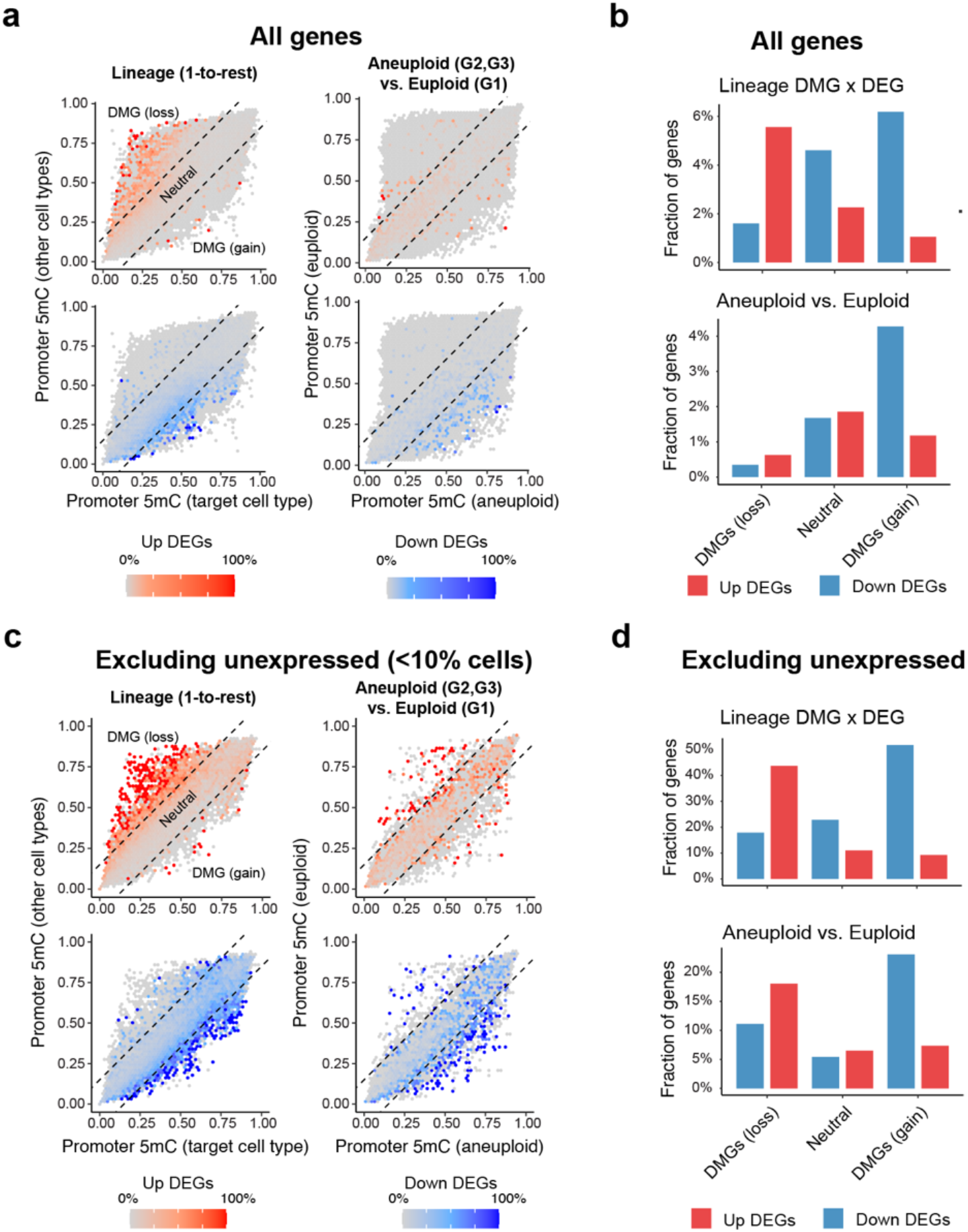
Differential DNA methylation mediate transcriptional changes in cell lineage establishment and disease progression. **a**, Relationship between promoter DNA methylation levels (x- and y-axis) and corresponding differential gene expression (color scale). The left panel compares promoter 5mC levels in a given cell type (x axis) versus other cell types (y axis), whereas the right panel compares aneuploid luminal epithelial cells from G2 or G3 (x axis) to euploid luminal epithelial cells from G1 (y axis). DEGs that are upregulated are shown in red and downregulated DEGs in blue. Dashed lines indicate ±15% differences in 5mC levels in the target cell type or sample group, corresponding to hyper- or hypomethylation relative to the reference. **b**, Fraction of DMGs that overlap with DEGs. **c-d**, Same analyses as in **a** and **b**, except that the gene set was prefiltered to exclude lowly expressed genes, defined as those detected in fewer than 10% of cells in either the target or the reference population in DEG analysis.

**Supplementary Figure 9.**
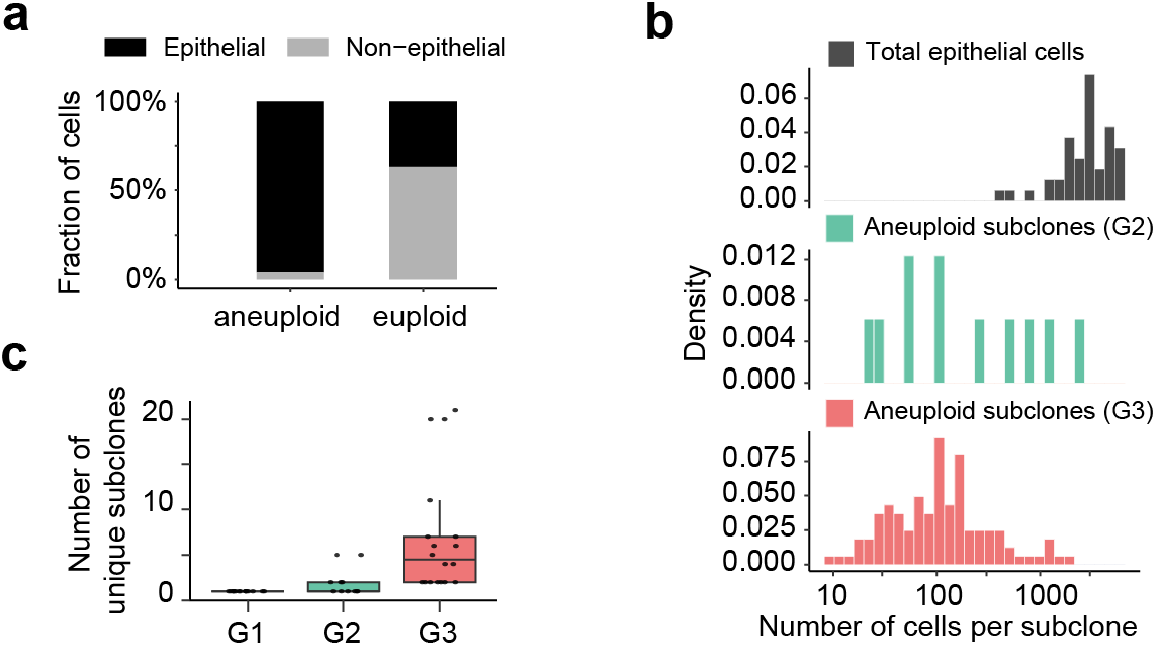
Composition of CNV-altered cells in prostate cancer. **a**, Proportion of aneuploid and euploid cells across all cells from 44 patient samples. **b**, Subclonal size defined as the number of cells sharing an identical karyotype forming each patient-derived subclone in G2 or G3. The top panel shows the total number of epithelial cells per patient. **c**, Distribution of subclone numbers across G1-G3. Euploid genomes were counted as a single subclone.

**Supplementary Figure 10.**
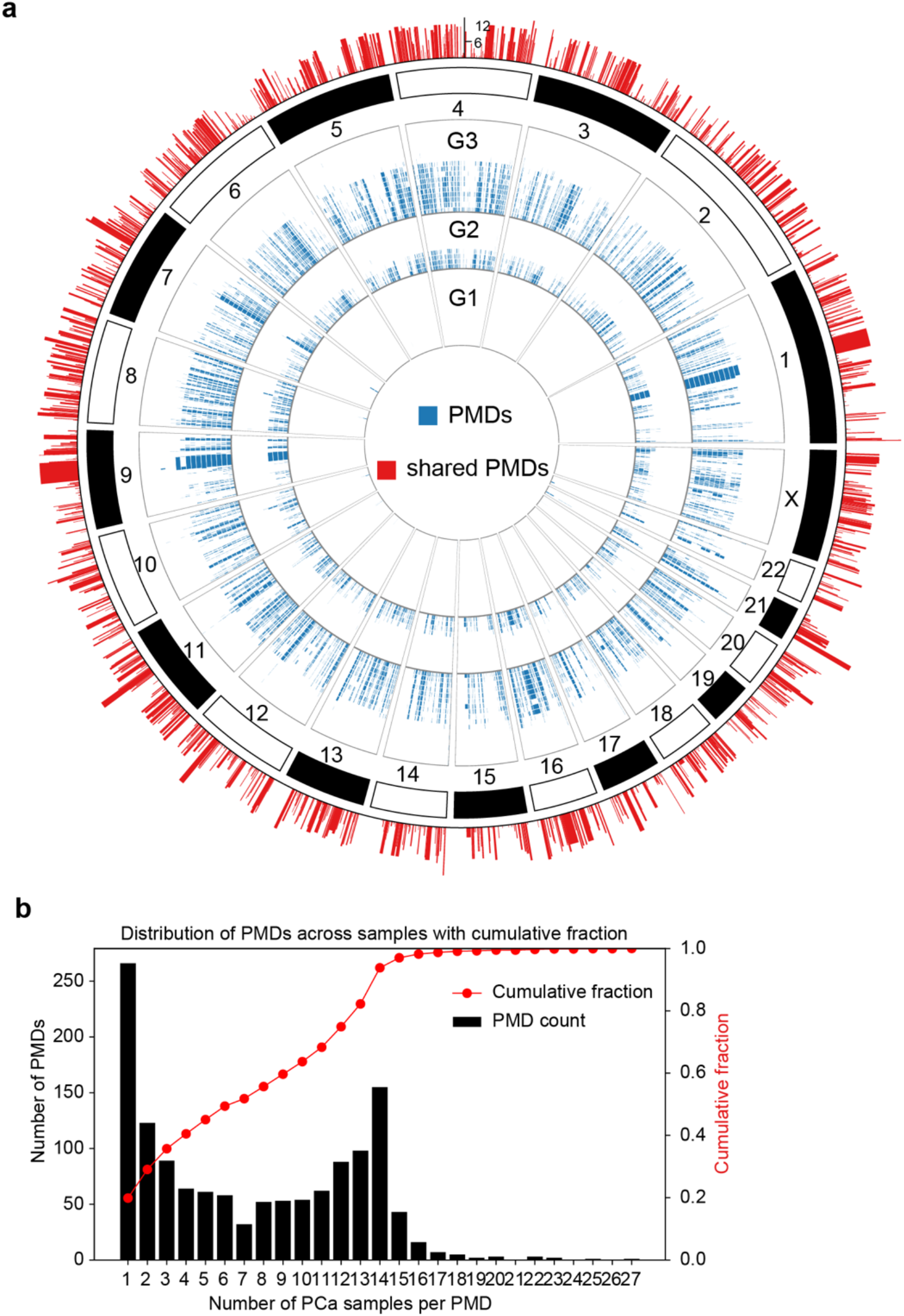
Identification of partially methylated domains (PMDs). **a**, Circos plot showing the genomic distribution of PMDs across 44 samples. PMDs shared between samples are represented as a frequency bar plot in the outer ring. **b**, Histogram of PMDs shared across multiple PCa samples. The overlaid curve shows cumulative fraction of PMD shared by *N* samples.

**Supplementary Figure 11.**
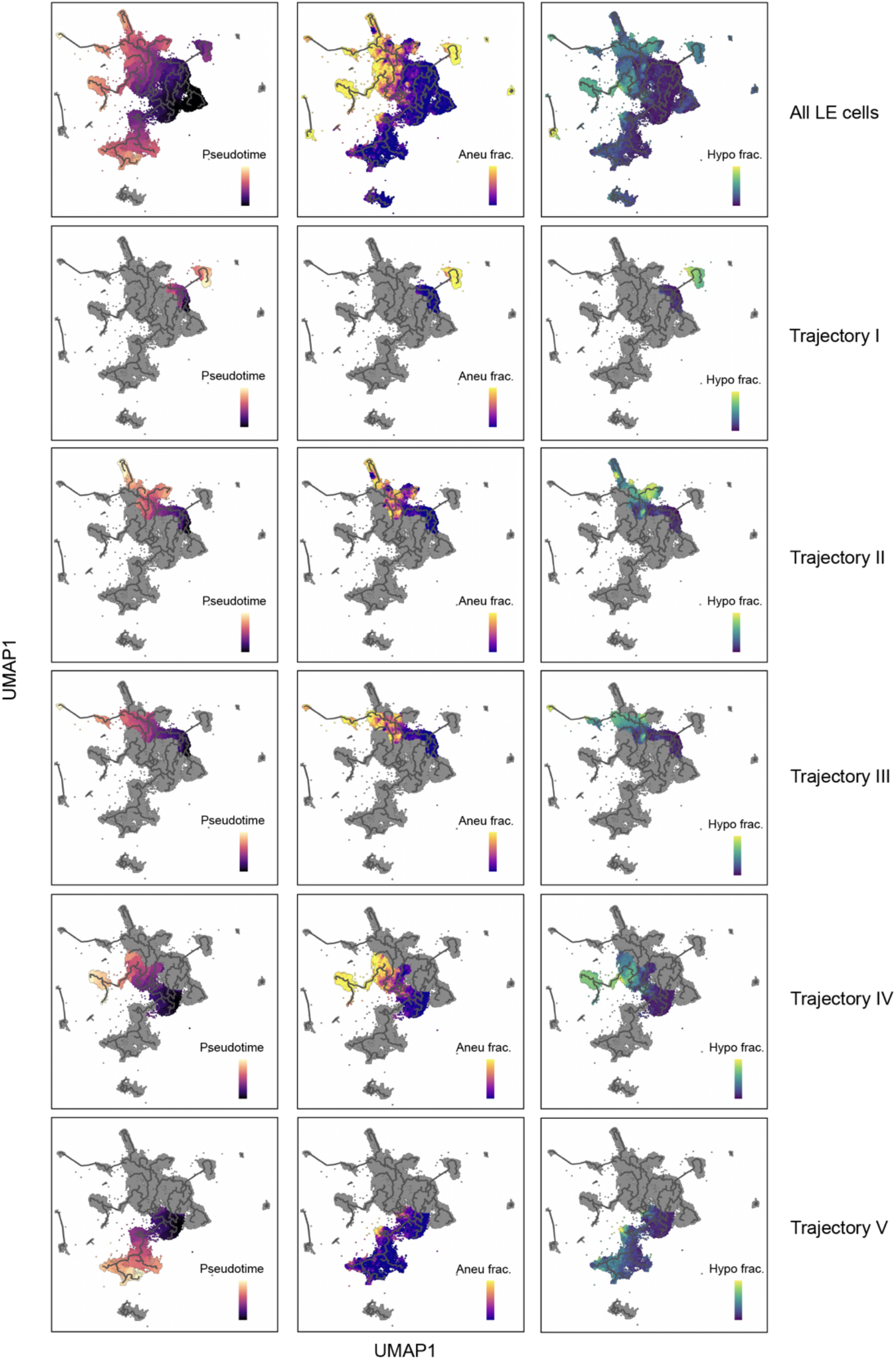
Transcriptome-centric pseudotime trajectory of epithelial cells. UMAP embedding and pseudotime were generated using Monocle3 on luminal epithelial cells from 44 patients. The root cluster was manually assigned after inspection of Leiden clusters in UMAP space, selecting a cluster devoid of aneuploid cells and spatially distant from aneuploid populations. The inferred Monocle3 trajectory graph is overlaid as lines across cell clusters.

**Supplementary Figure 12.**
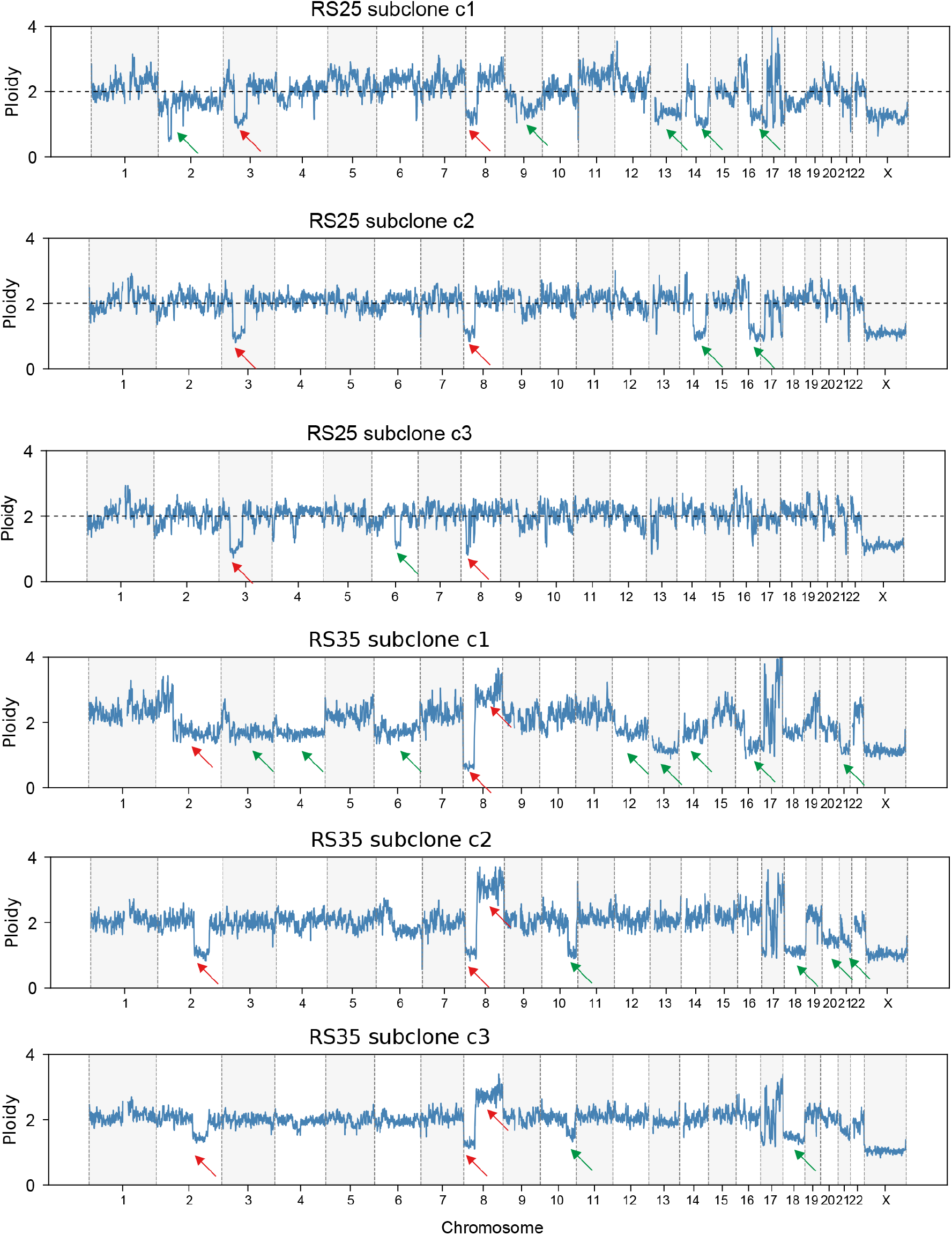
Refined CNV profiling of subclonal karyotype in sample RS25 and RS35. DNA sequencing reads from LE cells assigned to aneuploid RS25 and RS35 subclones were merged into pseudo-bulks. Randomly downsampled euploid cells were used as controls for normalization of read coverage. Arrows denote consensus CNV events in each subclone (Red: shared CNVs; Green: lineage-specific CNV).

**Supplementary Figure 13.**
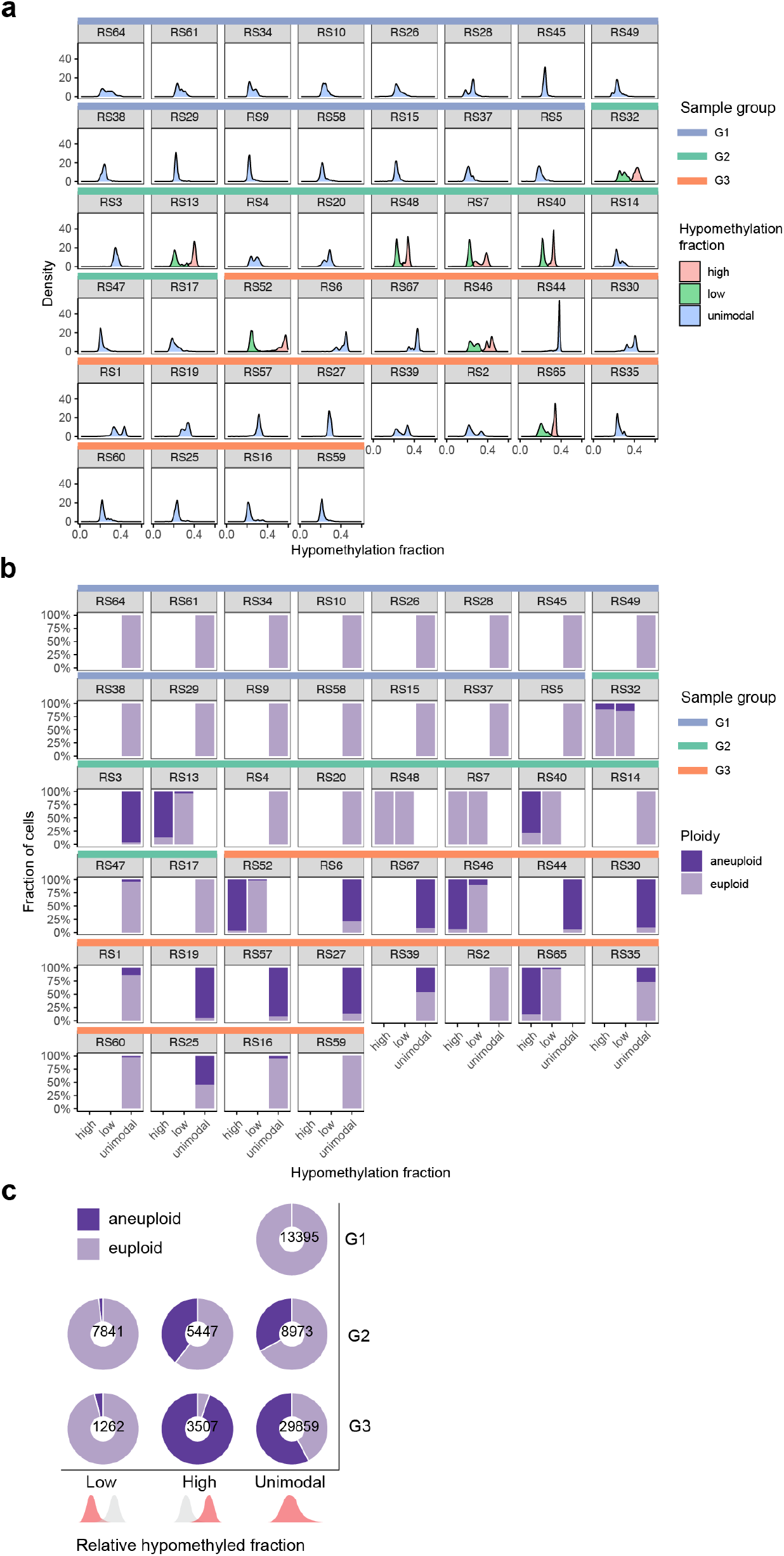
Bimodal distribution of global hypomethylation fractions in PCa samples. **a**, Distribution of hypomethylation fraction per LE cell, fitted using a Gaussian mixture model (G = 1-2) to determine whether the distribution is unimodal or bimodal. In samples exhibiting bimodality, cells were classified into low and high hypomethylation subpopulations **b**, Composition of LE populations (high, low and unimodal) in relation to aneuploidy fraction. **c**, Summary of aneuploid vs. euploid proportions across low, high and unimodal populations.

**Supplementary Figure 14.**
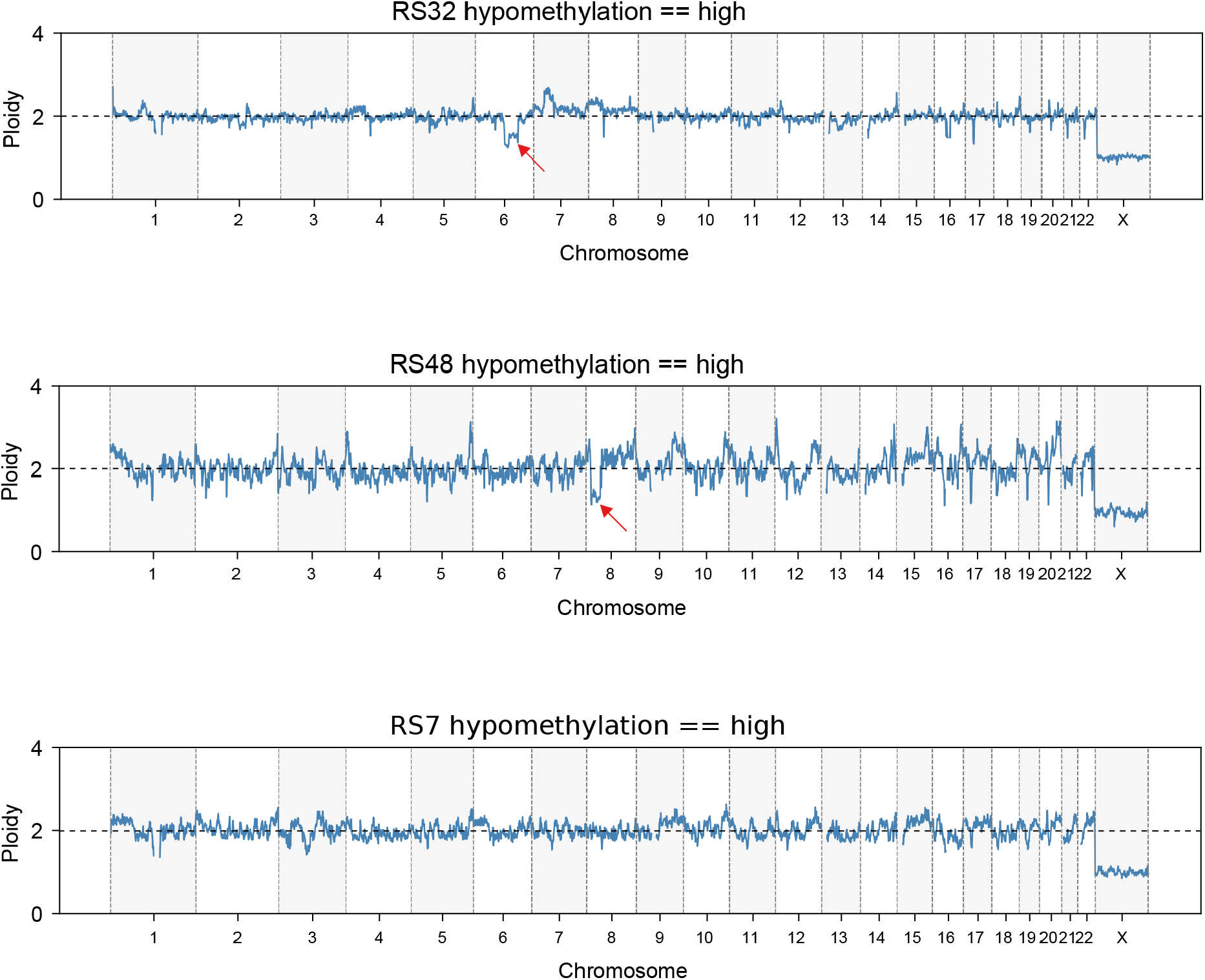
Verification of chromosomal integrity by aggregating single-cell DNA sequencing reads in pseudo-bulk. DNA sequencing reads from three samples (RS32, RS48 and RS7) were aggregated for LE cells with high hypomethylation fraction to generate pseudo-bulks, which were then inspected for CNV across the genome. Read coverage was normalized of each 250kb genomic bin in the target population by that of a reference population randomly downsampled from euploid cells. The normalized coverage was subsequently scaled by a factor of 2 to represent diploid copy number, with the exception of chromosome X treated as haploid. Arrows denote the position of consensus CNVs in the examined cell population.

**Supplementary Figure 15.**
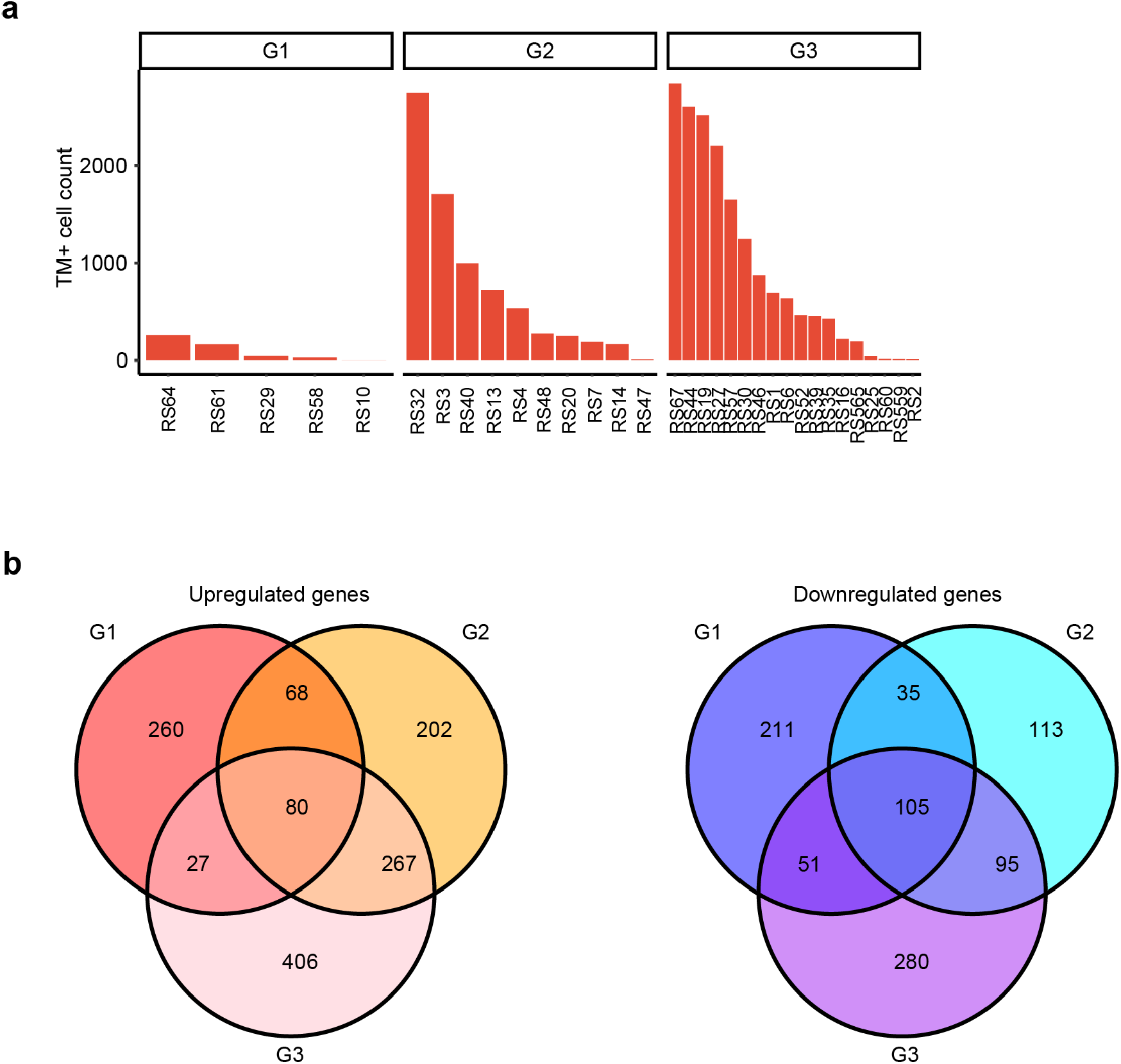
DEG analysis for TM^+^ cells relative to normal euploid LE cells. **a**, TM*^+^* cell count distribution across patients. Samples without qualified TM*^+^*cells were disregarded. **b**, Venn plots showing overlap of DEGs between euploid TM*^+^* and TM*^−^* cells (normal LE cells) among three groups.

**Supplementary Figure 16.**
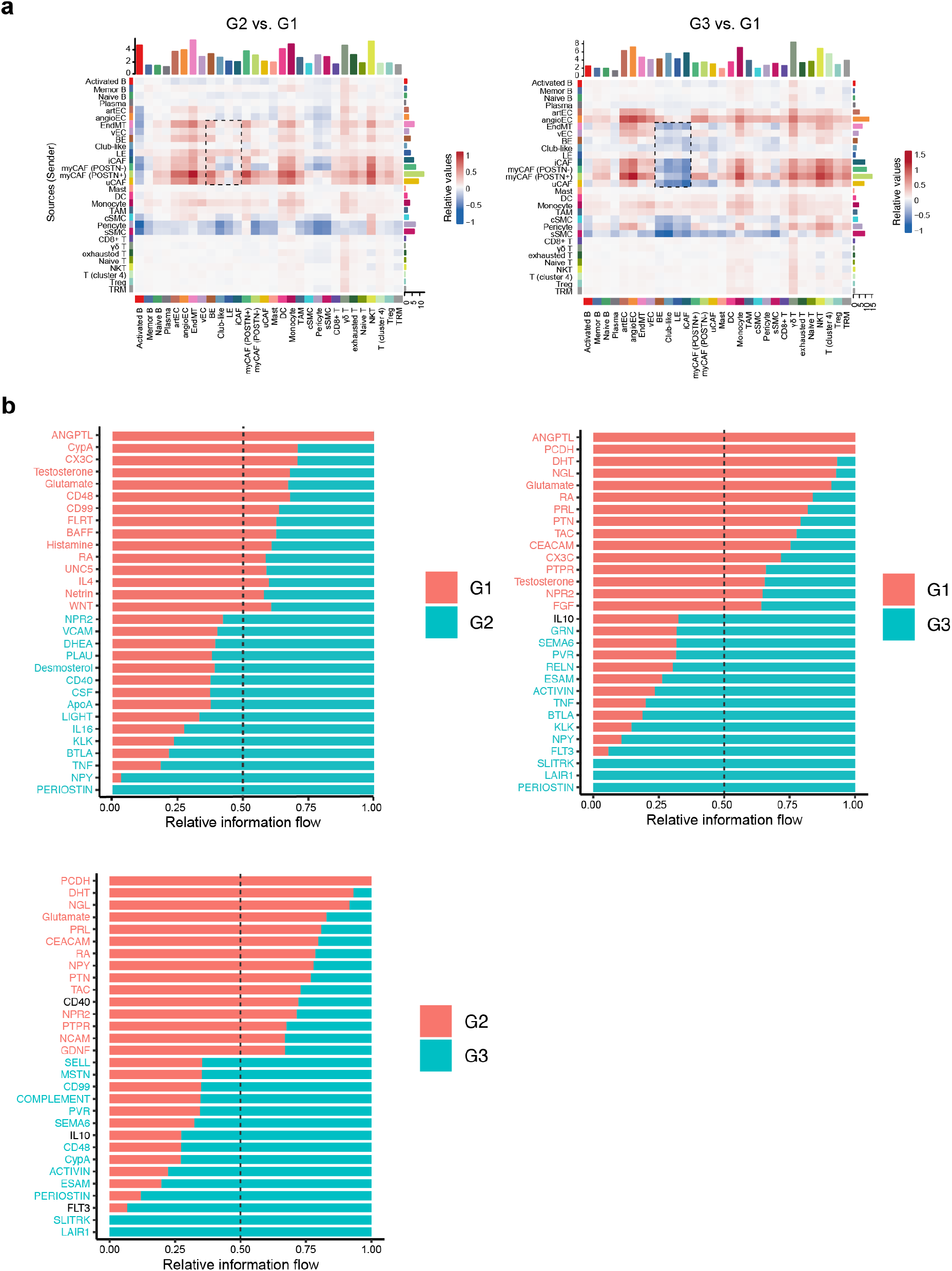
CellChat analysis of differentially expressed signaling pathways. **a**, CellChat analysis for 30 cell types in PCa (G2, G3) relative to benign controls (G1). Heatmap shows signaling strength. **b,** Pairwise comparisons include all cell types; only the top 10 upregulated and downregulated pathways in the second group relative to the first are shown.

